# Super-resolving particle diffusion heterogeneity in porous hydrogels via high-speed 3D active-feedback single-particle tracking microscopy

**DOI:** 10.1101/2025.03.13.643103

**Authors:** Yuxin Lin, Haoting Lin, Kevin D. Welsher

## Abstract

Nanoparticle diffusion in 3D porous structures is critical to understanding natural and synthetic systems but remains underexplored due to limitations in traditional microscopy methods. Here, we use 3D Single-Molecule Active-feedback Real-time Tracking (3D-SMART) microscopy to resolve nanoparticle dynamics in agarose gels with unprecedented spatiotemporal resolution. We highlight ‘hopping diffusion’, where particles intermittently escape confinement pockets, providing insights into hydrogel microstructure. Long, highly sampled trajectories enable extraction of kinetic parameters, confinement sizes, and thermodynamic barriers. This study demonstrates 3D-SMART’s ability to probe particle-environment interactions at super-resolution (∼10 nm in XY and ∼30 nm in Z) in 3D, offering new perspectives on nanoparticle diffusion and the structural dynamics of porous materials, with implications for drug delivery, material science, and biological systems.

**ToC figure:** 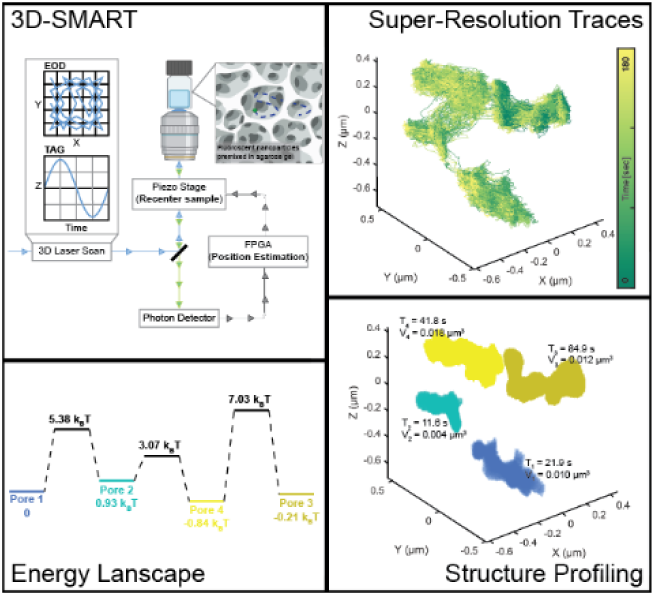

## 1. Introduction

In recent years, the development and application of single-particle tracking across many fields has been widespread.^[1–4]^ However, nanoparticle (NP) diffusion and interaction within 3D structures remain underexplored. Nanoparticle diffusion within porous microstructures is scientifically intriguing and crucial in both natural and synthetic systems. Porous microstructures dramatically impact the diffusion and penetration of nanoparticles. For example, mucus clearance acts as a primary barrier against viral infection and serves as an obstacle to be overcome for intranasal drug delivery.^[5, 6]^ In gel electrophoresis and column chromatography, the diffusion of nanoparticles within complex 3D environments may provide critical structural characterization.^[7]^ In porous biological and synthetic materials, nanoparticle transport is often heterogeneous due to variations in pore size and microenvironments.^[8–10]^ Such diffusion heterogeneity directly influences processes ranging from drug delivery and virus transport through mucus to the mechanical and structural characterization of biomaterials. Resolving this heterogeneity is essential for understanding nanoparticle-matrix interactions, optimizing delivery strategies, and engineering functional materials with predictable transport properties. However, characterizing these spatially and temporally varying diffusion behaviors at the nanoscale within three-dimensional porous matrices remains challenging.

Electron and atomic force microscopies are common tools for studying the structure and mechanical properties of hydrogels and other microporous structures. Unfortunately, both typically involve perturbative experimental requirements that irreversibly alter the microstructures and cannot resolve particle dynamics within the hydrogel.^[11, 12]^ Single-particle tracking (SPT) microscopy, or microrheology, is a frequent tool for studying the interaction of nanoparticles and microstructures.^[10]^ Trajectories of single particles are usually achieved using image-based methods, in which the particle of interest is imaged sequentially, and localizations are concatenated into a trajectory.^[2, 13]^ However, limitations exist for image-based tracking methods in terms of observation duration, lack of spatiotemporal resolution, and restriction to two-dimensional motion.^[14–16]^ When particle motion is confined to two-dimensions (and just one focal plane), the temporal resolution is limited by the time it takes to collect one image, determine by the number of pixels and pixel dwell time for confocal methods, and camera exposure and readout for widefield methods. However, for three-dimensional processes, the particle localization rate is limited by the volumetric imaging rate, which is the single focal plane imaging rate multiplied by the number of planes required, dramatically decreasing the temporal resolution. One approach to break this volumetric imaging bottleneck is to apply point spread function (PSF) engineering to encode axial information into the 2D image of a single particle. While most PSF engineering approaches are restricted to an axial range of a few microns,^[17, 18]^ more recent PSF engineering techniques were able to extend the axial range to tens of microns with complex phase modulations and specially designed optics.^[19, 20]^ However, the scattering induced by the thick and complex 3D matrix will distort PSF patterns and pose a further challenge to PSF engineering methods. An alternative approach called fcsSOFI,^[7, 21]^ developed by Kisley *et al*, combines super-resolution optical fluctuation imaging (SOFI) and fluorescence correlation spectroscopy (FCS) and is a powerful tool to characterize spatial and diffusion information within porous materials. fcsSOFI overcomes the problem with limited sampling in SPT but sacrifices dynamic information that can be extracted from single particle trajectories. Also, in most applications of the methods mentioned above, single-particle trajectories are very challenging to understand due to high noise and short observation time, diminishing the potential to understand the behaviors of individual particles in detail. Short, noisy, and low sampling rate trajectories limit the knowledge that can be extracted. Often, the best that can be expected is the measurement of a diffusion coefficient or extraction of non-Brownian behavior from an ensemble mean-square displacement (MSD) analysis. Long, highly sampled, three-dimensional trajectories are needed to extract the structure and dynamics of porous materials. In this work, we demonstrate that active-feedback 3D single-particle tracking microscopy is well-suited for super-resolving gel structure and dynamics in three dimensions.

To date, the primary method for investigating diffusion heterogeneity in porous gel materials has been microrheology enabled by single-particle tracking. Previous reports have used either wide-field fluorescence microscopy,^[15, 16]^ or spinning disk confocal microscopy.^[14]^ In both methods, images can be collected at reasonable frame rates, but with only two-dimensional information. This means that a particle may appear stationary, but may actually be rapidly moving along an unseen dimension. Confocal approaches can, in theory, address this shortcoming. However, scaling a single confocal frame into a volume dramatically reduces the temporal resolution to a point that the trajectories do not yield useful information on their dynamics or the local environment. Furthermore, even in two-dimensions, the available temporal resolution (10-50 Hz) is insufficient to capture the structure of the pores within these gels, and have little to no chance to measure particle dynamics within the narrow channels between pores. Clearly, a faster, three-dimensional SPT method is needed to detail this complex diffusion process.

Particle diffusion within porous materials is often treated as anomalous diffusion, typically confined motion due to the physical hindrance. Typical analytical approaches used include mean squared displacement (MSD) analysis,^[22]^ change-point detection,^[23]^ hidden Markov model.^[24]^ These methods have proven their effectiveness in probing anomalous diffusion, primarily by examining how diffusion behavior evolves over time. However, as mentioned above, the diffusion heterogeneity within porous materials fundamentally originates from spatial structural heterogeneity. In this complex scenario, conventional analytical methods can average over spatially distinct micro-environments, leading to a less sensitive and accurate characterization. For particle diffusion within narrow cavities, rapid sampling over extended periods is necessary to map both the particle dynamics and the local structure. To achieve such dense sampling in nanoscale environments, highly sampled trajectories from active-feedback tracking methods are required.

Recent years have seen a wide range of applications of active-feedback tracking microscopy.^[3, 25–29]^ Active-feedback microscopy presents a unique opportunity to demystify hydrogel microstructures along larger spatial, especially axial, scales with greater detail. Instead of taking images sequentially, active-feedback tracking microscopy uses modified detection^[30]^ or patterned excitation^[31, 32]^ to obtain real-time localization information of rapidly diffusing particles and uses a closed feedback loop to “lock” the particle in the center of the observation volume. Active-feedback tracking microscopy uses previous particle position and updated photon information for real-time location estimation; thus, by using information from prior localizations, fewer photons are required for better spatial and temporal resolution compared to image-based methods. Also, the typical excitation or detection module is three-dimensional. It is not required to take images at multiple planes and assemble them into a stack (z-stacking) for tracking along the axial direction, which greatly enhances the sampling rate for 3D tracking. Similar to active-feedback microscopy, PSF engineering techniques can also achieve high volumetric sampling rates by encoding axial information into a single 2D image. However, these approaches typically operate in widefield or epifluorescence illumination modes, where the entire sample volume is illuminated. In scattering media such as hydrogels, this broad illumination can result in increased PSF distortion due to scattered excitation and emission photons. In contrast, most active-feedback tracking methods utilize confocal or point-scanning architectures, which restrict excitation and detection to a small focal volume, thereby reducing PSF distortion and enhancing robustness against scattering-induced degradation.

As one of the active-feedbacking tracking methods, 3D Single-Molecule Active-feedback Real-time Tracking (3D-SMART) microscopy has been employed to monitor polymer growth,^[33]^ visualize viral infection,^[34–36]^ and screen lipid nanoparticle formulations for enhanced delivery of mRNA in live cells and for optimal mucus penetration^[37, 38]^. This broad application scope makes 3D-SMART a good candidate to resolve the highly heterogenous diffusion of particles within porous scaffolds.

Here, we used 3D Single-Molecule Active-feedback Real-time Tracking (3D-SMART) microscopy to monitor the three-dimensional dynamics of nanoparticles within agarose gels (AG) with high spatiotemporal resolution and long tracking duration. Notably, in larger nanoparticles and higher concentration AG preparations, it is frequently observed that particles can repeatedly “hop” between confinement cages or explore the interior of hydrogel in a hopping diffusion manner. Kinetic parameters, such as the lifetimes, on/off rates, and hopping energy barriers can be extracted from highly sampled trajectories. The highly sampled trajectories depicted the complex 3D environment and could be utilized to probe the microstructures with super-resolution of ∼ 10 nm.

## 2. Results

### 2.1. High spatiotemporal SPT via 3D-SMART

Agarose gels (AG) were chosen as the model system for this work due to their extensive use in biomolecule separations and thorough characterization.^[39]^ AG also has a refractive index close to aqueous buffer and is macroscopically uniform;^[40]^ therefore, severe aberrations would not be expected at the tracking depths used in this work. Fluorescent nanoparticles of different sizes were premixed within agarose gels of desired concentrations. High-speed single-particle tracking was accomplished by 3D-SMART microscopy as previously described with no required modification (**Figure 1**, **Figure S1**).^[31, 34, 41]^ Briefly, a 3D scanning volume of 1 × 1 × 2 μm in XYZ was generated laterally with a pair of electric-optical deflectors (EODs, ConOptics, M310A) and axially with a tunable acoustic gradient (TAG, TAG Optics, TAG 2.0) lens.^[42]^ The lateral plan is a 5 × 5 grid sampled at 20 µsec per spot, while the focus is simultaneously scanned at 70 kHz in Z. The photon arrival information was captured by a single photon counting avalanche photodiode (APD, Excelitas, SPCM-ARQH-25) and applied to estimate the real-time position of the particle with an assumed Gaussian density Kalman filter.^[43]^ The real-time position estimation was used to drive the piezoelectric stage to ‘lock’ the diffusing particle in the center of the scanning volume. The average excitation power of the 488-nm beam at the focus was 45-180 nW for 200-nm probes, 90 nW for 100-nm probes, and 9 nW for 500-nm probes. In theory, different EOD scan patterns and smaller scan sizes could increase tracking precision at the cost of the ability to track fast-diffusing particles.^[34, 44]^ For the benefit of tracking robustness and consistency across different experimental conditions, parameters were chosen that allowed tracking of particles free in solution, as well as within porous hydrogels.

**Figure 1.**
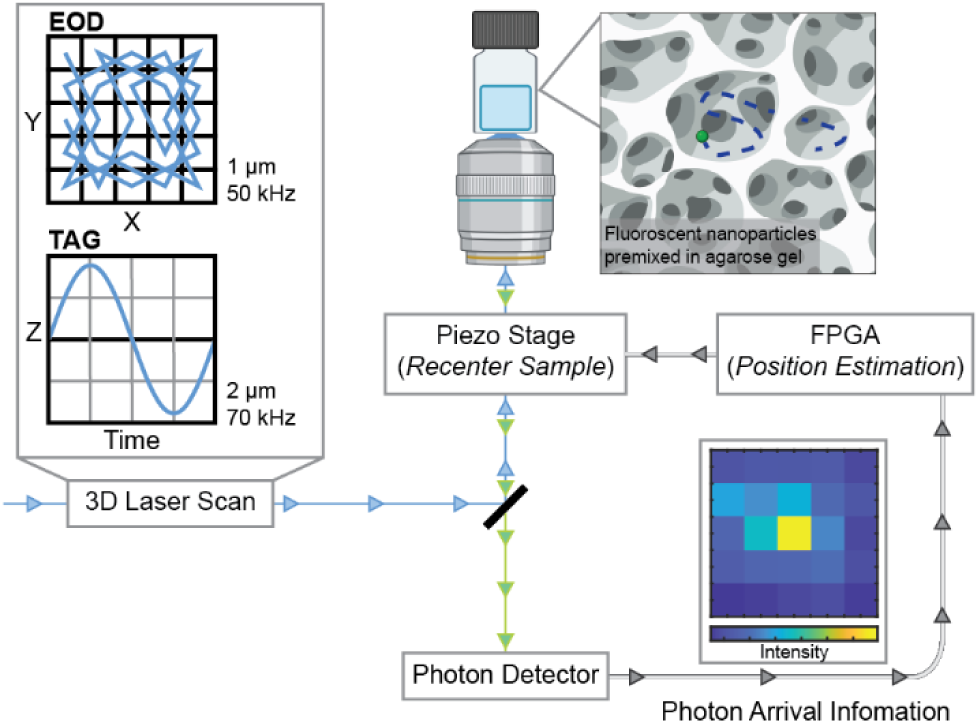
Illustration of performing SPT in hydrogels using 3D-SMART. A sub-millisecond 3D scanning volume of 1×1×2 μm is generated by a pair of EODs and a TAG lens. Probes were premixed with the desired concentration of agarose gel solution and solidified in the chamber. The photon detector (APD) collects the emitted photons, and photon arrival information is routed into an FPGA for real-time position estimation of the diffusing probe. The updated localization information is used to drive the piezoelectric stage to effectively hold the diffusing nanoparticle in the center of the laser scan.

The position sampling rate of 3D-SMART is 1 kHz, corresponding to the piezoelectric stage’s response time. This sampling rate can be improved by using feedback elements with faster response times. In prior work, a galvo scanning mirror was employed to replace the XY piezoelectric stage, which improved the sampling rate to approximately 5 kHz.^[25]^ The sampling rate can theoretically be further enhanced with brighter probes.^[45, 46]^ Since a single-photon counting APD is used for detection, there is no exposure or readout time limitation. The particle position can theoretically be updated with each photon.

Previous work from our group has validated that the tracking algorithm retains reliable localization accuracy in environments with moderate background fluctuations.^[47, 48]^ AG-induced PSF aberrations primarily broaden the PSF or introduce mild asymmetry.^[49]^ In this case, the active-feedback nature of 3D-SMART continuously updates the estimated centroid positions in real time, effectively compensating for mild PSF asymmetry and mitigating its impact on tracking performance. However, in scenarios where significant centroid shifts occur due to severe aberrations, tracking accuracy would inevitably be compromised. In practice, the lateral localization precisions are about 10 nm with 300 kHz intensity (i.e., 300 photons in 1 ms bin time, **Figure S2**a-b). Compared to non-gel tracking, the axial tracking localization precision (29.7 ± 2.8 nm) in AG was slightly reduced due to scattering in the gel network (Figure S2c).

In addition to the 1 kHz position sampling of 3D-SMART, the photon arrivals can be assembled into an image called the “EOD Image.” The EOD image is a critical piece of information that allows confirmation that trajectories arise from a single particle (**Figure S3**, also see Methods section for more details).

### 2.2. Particle diffusion is hindered by concentrated hydrogel and large particle size

First, passive diffusion of fluorescent nanoparticles of various sizes within the agarose gel of different concentrations was observed (see Methods). The results and tendencies were very intuitive and similar to expected and previously reported results.^[8, 39]^ For probes of the same size, the degree of spatial confinement increased with increasing agarose concentration, leading to more hindrance of the particle diffusion (**Figure 2**A, **Figure S6**a-c). While for the same agarose gel, the larger beads tend to be more confined while smaller beads diffuse freely (Figure 2B, Figure S6d-f). Trajectories could be quantified using diffusion coefficient (*D*), power exponent *α*, and packing coefficient (*Pc*).^[50]^ Please refer to Supplementary Information for details.

**Figure 2.**
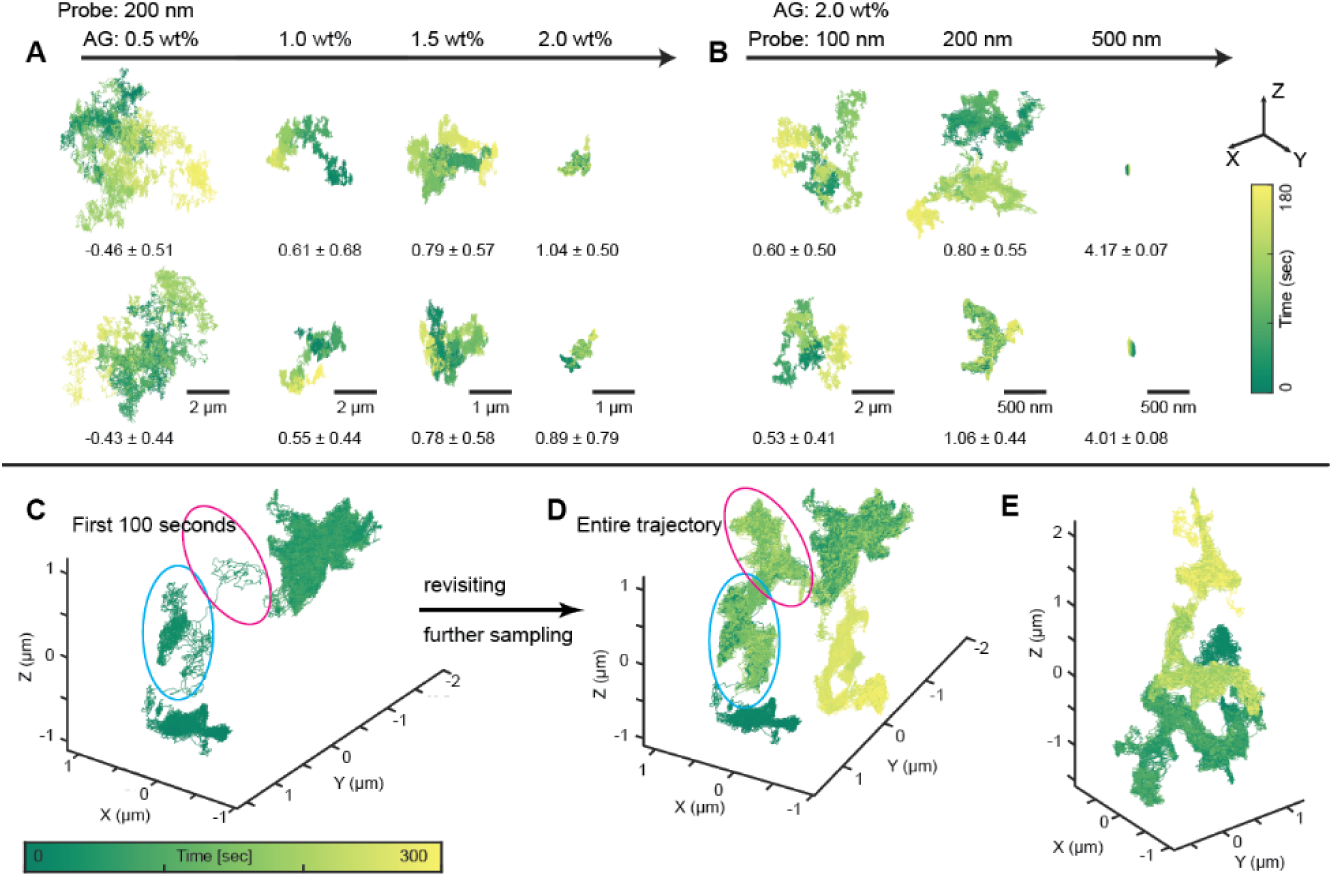
Mesh-size- and probe-size dependent diffusion. (A) Representative trajectories of 200 nm probes diffusing in increasing wt% AG (see detailed information in Figure S4. N = 51 and 31 for control groups of free and fixed respectively. N = 59, 52, 127, and 93 for 0.5, 1.0, 1.5 and 2.0 wt% AG, respectively). Scale bars: 0.5 wt% and 1.0 wt%: 2 μm; 1.5 wt% and 2.0 wt%: 1 μm. (B) Representative trajectories of probes of different sizes diffusing within 2.0 wt% AG (see detailed information in Figure S5. N = 141, 127, and 57 for 100, 200, and 500 nm probes, respectively). Scale bars: 100 nm probes: 2 μm; 200 nm and 500 nm probes: 500 nm. All trajectories in (A) and (B) share the same color bar and axis. Calculated results of packing coefficient (mean ± std in one second segments) are listed below each trajectory. (C and D) the first 100-second segment and the entire trajectory of a 200 nm probe diffusing within 2 wt% AG. The red and blue circles highlight two cavities were initially bypassed but eventually revisited after multiple hopping events. (E) Another example of a 200 nm probe diffusing within 2 wt% AG covering a relatively large 3D volume. The same color bar from (C) applies to all three plots (C-E).

Presumably, as the particle size approaches the pore size, particles exhibit more confined diffusion. In some cases, probes dwelled in or repeatedly visited the same confinement pore. For 200 nm probes, when the AG concentration reached 1.0 wt%, hopping diffusion started to be observed: particles underwent confined diffusion in one area and intermittently escaped to neighboring areas through connective channels. Hopping escapes between two confinements always featured larger displacement and were highly one-dimensional, in contrast to confined diffusion (**Figure S7**b-c). Various diffusive behaviors were observed after hopping events. Some particles hopped to relatively free diffusion (see examples in **Figure S4**b), while others hopped to a neighboring pore and exhibited confined diffusion (see examples in Figure S4c-d). With the combination of larger particles and denser AG, e.g., 2 wt% AG and 200 nm probes, hopping diffusion became a major diffusion pattern for particles to explore the porous hydrogel passively.

Due to the 3D nature of hydrogels, the ability to resolve diffusion and confinement structures in 3D is critical. In active feedback tracking microscopy, particles can be tracked across an ample axial range without sacrificing spatial or temporal resolution. An example of 3D pore-to-pore hopping diffusion is depicted in Figure 2C-D (see also **Figure S8**a-b and **Movie S1**), in which the particle hopped through four major cavities in a nonconsecutive manner. After being trapped in the first pore for ∼ 25 seconds, the particle bypassed two cavities briefly and thoroughly explored the fourth cavity first. However, the particle swung back eventually, further sampling the second and third previously ignored confinements. Another example trajectory (Figure 2E, see also Figure S8c-d and **Movie S2**) shows that hopping diffusion also occurs across a wide axial range, posing challenges to conventional tracking methods to capture and resolve. The ability to track and resolve 3D spatial information is critical when tracking smaller probes in looser meshes due to increased diffusivity.

### 2.3. Thermodynamics of hopping diffusion

The high sampling rate and precise localization of 3D-SMART enabled analysis of the thermodynamics and kinetics of hopping diffusion and characterization of 3D structure in AG beyond what is possible with traditional SPT. Extraction of the 3D nanoscale structure is required to determine the underlying thermodynamic within heterogeneous and porous materials. To that end, confinement pockets of diffusing particles were identified and segmented based on the dwell time in each pixel (2D) or voxel (3D) (see Methods for details). For example, as shown in **Figure 3**, a 180-second 3D trajectory of a 200 nm probe diffusing within 2.0 wt% AG (Figure 3A) was binned into 10×10×20 nm voxels (XYZ, determined by 3D localization precision). This 3D binned data was then converted into a 3D heatmap representing the dwell time within each voxel. The 3D hotspots in the heatmap correspond to the confinement pockets where the particle repeatedly visited and dwelled. Based on the intensity of the heatmap, the voxels were grouped, allowing the identification of four distinct confinements of different shapes and sizes (Figure 3B). The volume of the confinement was then calculated from the number of voxels, while the dwell time for each confinement was calculated from the number of localizations within a confinement.

**Figure 3.**
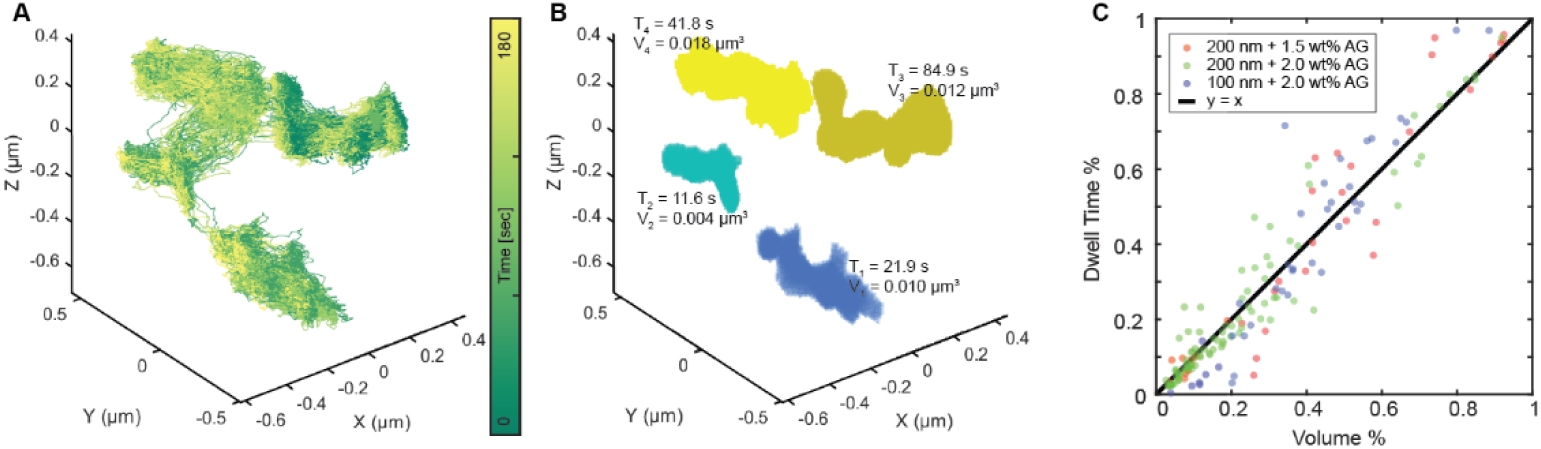
(A) 3D trajectory of a 200 nm probe’s hopping diffusion within 2.0 wt% AG. Trajectory is color-coded by time. (B) Four identified confinements of the trajectory shown in (A), with corresponding dwell time (T) and confinement volume size (V). The color code represents different confinements. (C) Plot of the fractional volume against the fractional dwell time, indicating hopping diffusion is driven by entropic differences between confinement areas.

To better understand the thermodynamics of hopping diffusion, we analyzed the relationship between fractional dwell time and fractional confinement volume. Fractional dwell time was calculated as the dwell time in each confinement divided by the total dwell time across all confinements within the trajectory. Fractional confinement volume was similarly defined as the volume of each confinement divided by the total volume of all confinements. Three experimental setups were analyzed: 200 nm probes in 1.5 wt% AG, 200 nm probes in 2.0 wt% AG, and 100 nm probes in 2.0 wt% AG. In all cases, a clear one-to-one relationship between fractional dwell time and confinement volume was observed (Figure 3C), indicating that hopping diffusion is governed purely by entropic differences between the pores (see Methods for details), which is determined by pore size. Furthermore, this relationship suggests that the free energy difference, Δ*F*, between two adjacent confinements is proportional to the ratio of their confinement volumes:

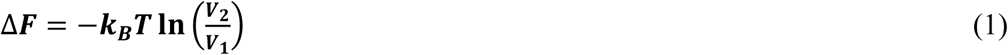

where *k*_*B*_ is the Boltzmann’s constant, *T* is the absolute temperature, in this manuscript *k*_*B*_*T* is used as a unit of energy, *V* is the volume of a specific confinement during hopping diffusion.

### 2.4. Kinetic information extracted from hopping events

Although there exist some prior experimental observations of hopping diffusion, kinetic and thermodynamic information, such as the free energy barrier, remains very challenging to extract due to the limitation in spatiotemporal resolution.^[16, 51, 52]^ Moreover, multiple factors, including the particle diameter, size of the channel, and length of agarose polymers occupying the channel, may determine the free energy barrier the particle needs to hop from one pore to another.^[53, 54]^ The heterogeneous nature of the agarose gel makes it challenging to probe the energy barrier from hopping diffusion in typical single-particle trajectories. Trajectories collected by 3D-SMART have the advantage of having high spatiotemporal precision, enabling frequently sampled and precise trajectories that can map out the entire environment of the particle, given a suitable duration of observation. Using 3D-SMART, it was observed that particles could hop between two adjacent pores repeatedly, allowing for a thorough sampling of the channel between the pores (**Figure 4**, see also **Movie S3**). The tracking performance during hops was examined to exclude artifacts (**Figure S9**c). From these trajectories, it is possible to gain a detailed understanding of the geometry of both the pores and the connecting channels and enable a detailed analysis of the kinetics of hopping diffusion.

**Figure 4.**
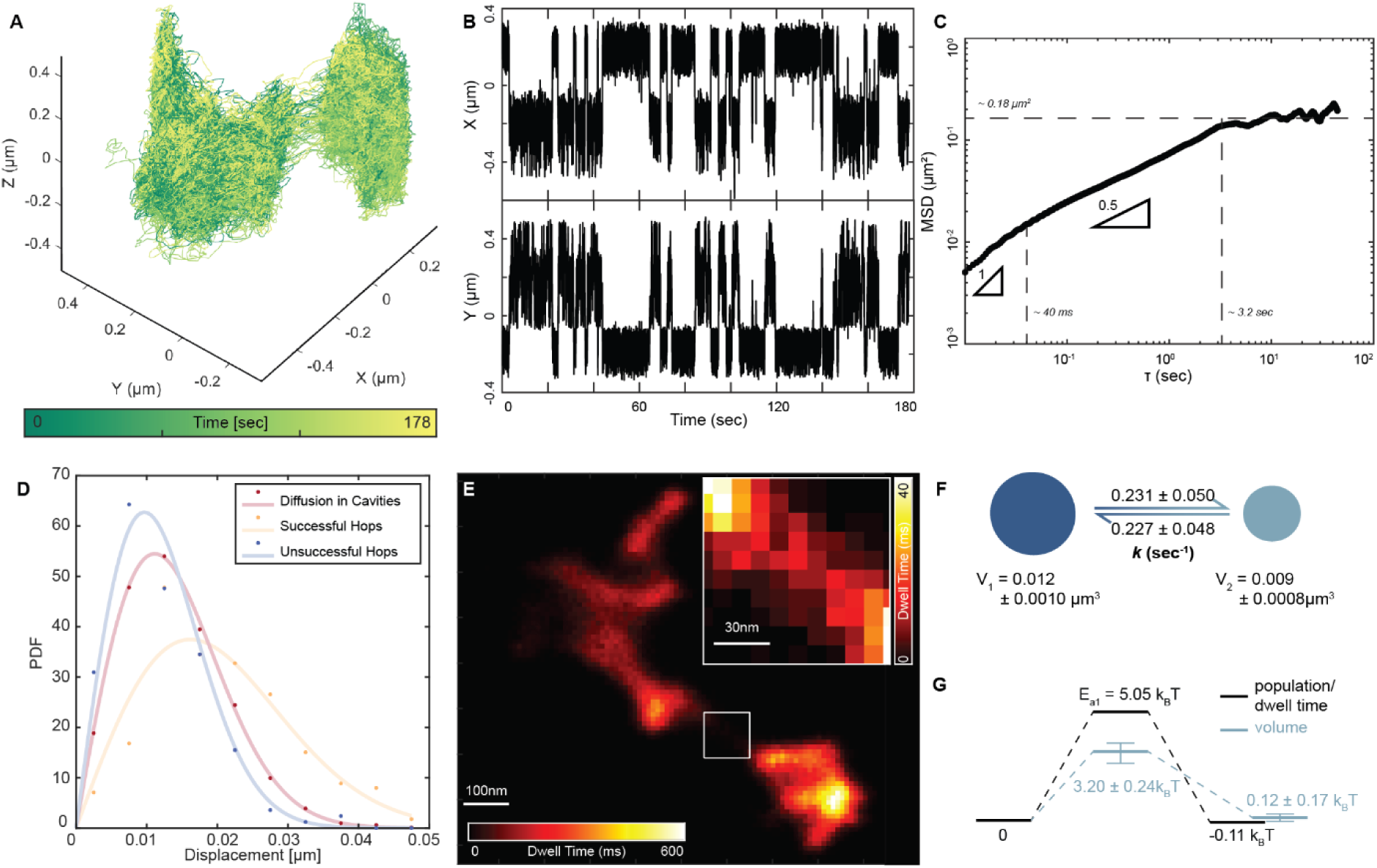
Illustration of a hop event. (A) Representative trajectory of hopping events between two cavities. NP size d = 200 nm, AG concentration c = 1.5 wt%. (B) X and Y traces of the trajectory shown in (A), indicating there existed two states for particle diffusion. (C) 2D MSD plot of the representative trajectory shows different diffusive states in cavities connected by a channel (minimum lag time of 10 ms). The two slopes were calculated based on a linear fit within the corresponding time window. Lag times shorter than 10 ms were chopped. see 3D MSD plot in Figure S10. (D) 100ms-Displacement distribution of diffusion in cavities, successful hops, and unsuccessful hops, respectively. Distributions were overlayed with Rayleigh distribution fitting. (E) 2D heatmap converted from 3D trajectory based on the number of localizations (or dwell time) within each pixel (10×10 nm). Inset: a zoomed-in view of the channel connecting two cavities. (F) Translocation rates, volumes, and areas of two confinements. (G) Energy diagram of the energy barrier between two confinements and the activation energy of the hop. The error bars in free energy barrier (calculated from volume) are determined by expanding or shrinking the channel volume by one localization precision.

To better analyze these hopping events, the confinement pockets of each 3D trajectory were identified using the algorithm mentioned above in both 2D and 3D. A 2D heatmap (density of the probe centroid localizations) is shown in Figure 4E and also serves as an indirect visualization of the interior microstructures carved out by the particle trajectory. Transitional localizations between two cavities were specifically extracted to measure the width of the “channel” connecting two pores and to measure the kinetics and thermodynamics of hopping events through these channels. The channel shown in Figure 4E is very narrow and only 68.0 ± 3.3 nm (Calculated as 3 × std representing ∼86% of the datapoints, Figure S9a-b) wider than the probe. Based on the intensity on the heatmap, the localizations of two states can be labeled accordingly as pores 1 and 2, with volumes of 0.012 ± 0.0010 and 0.009 ± 0.0008 μm^3^ (2D area size: 0.092 ± 0.0002 μm^2^ and 0.048 ± 0.0001 μm^2^, respectively (Figure 4F). Although their trajectories appeared relatively unremarkable along the axial dimension, the projected 2D area sizes of each confinement were 0.092 μm² and 0.048 μm², with a ratio of 1.9 (compared to the volume ratio of 1.3). This discrepancy may lead to an inaccurate estimation of the thermodynamic barrier. The translocation rate between the two confinements was also calculated as the inverse of lifetimes in each confinement (Figure 4F, Figure S9d-e), which were 0.231 ± 0.050 and 0.227 ± 0.048 s^-1^, respectively.

MSD analysis (2D MSD: Figure 4C; 3D MSD: **Figure S10**) reveals several different diffusion states for hop events. The particle first explored the cavities freely with exponent α ≈ 1 for the first ∼ 40 ms. After this free diffusion regime, the particle motion became subdiffusive with the MSD proportional to the square root of lag time up until ∼3 sec. After the probe thoroughly explored the pores, the MSD fluctuates around a maximum value 0.18 ± 0.02 μm^2^ (calculated based on MSD value after 3.2 sec), corresponding to a gyration diameter of 0.85 μm (0.94 μm for 3D). This gyration diameter provides a rough estimate of the area explored by the entire trajectory but may be less informative for irregularly shaped trajectories like the one presented here. Indeed, this value does roughly agree with the combined gyration diameters of the individual pores, which were ∼1.02 μm in 2D or ∼0.88 μm in 3D, respectively.

Hop events between pores are very short duration (∼ 6 ms, **Figure S7**a), highly one-dimensional along the channel (Figure S7b-c), and show slightly larger displacements (13 ^[8, 18]^ nm in pores, 19 ^[13, 27]^ nm channel, Median [Q1, Q3], Kruskal-Wallis Test, *p* < 0.001, Figure 4D). We further compared the displacement of unsuccessful hops, in which particles entered the channel but failed to pass through. These unsuccessful hops exhibited smaller displacements (11 ^[7, 16]^ nm, median [Q1, Q3], Kruskal-Wallis Test, *p* < 0.05) compared to diffusion in the pores (Figure 4D). This result suggests the presence of an entropic energy barrier that regulates hopping diffusion, and that larger displacements may facilitate successful translocation events. A more detailed discussion of this interpretation is provided in a later section.

Given the entropic nature of hopping diffusion and assuming thermal equilibrium over long timescale, the probability of finding a particle in each confinement follows Boltzmann distribution. This holds true not only for the two cavities but also for the connecting channel (Figure 4E), which was revisited over 40 times, yielding more than 300 data points. Therefore, the free energy difference between two confinements, Δ*F*, and the activation energy of hop events, *E*_*a*_, may be calculated based on the population (or occupancy/dwell time) within the cavities, respectively, as:

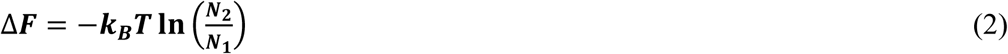

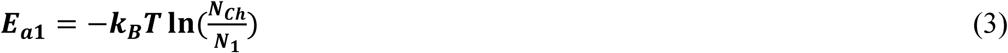

where *N_1_* and *N_2_* are the population of the two cavities, and *N_Ch_* is the population of the channel. *E*_*a*1_ is the activation energy barrier a particle needs to overcome to escape confinement 1.

The calculated free energy barrier to hop from pore 1 was 5.05 *k_B_T*, and the free energy difference between the two larger pores was -0.11 *k_B_T* (Figure 4G). To further validate the free energy barrier calculation, we examined its temporal dependence by computing the hopping free energy barrier through the channel as a function of time (Figure S14). The calculated free energy barrier stabilized after ∼ 60 seconds, indicating sufficient sampling and that the system had reached thermodynamic equilibrium. The free energy difference between the pores was also calculated based on the volumetric ratio using Equation 3, which gave a similar result (+0.12 ± 0.17 *k_B_T*, Figure 4G). The relatively small energy barrier between the two confinements explains the similar translocation rate between the two cavities.

It is worth emphasizing that hop events occur rapidly (∼6 ms), and cannot be observed by existing SPT microrheology methods. The population of trajectory points and the energy barrier for hopping diffusion can only be measured by methods with high spatiotemporal precision.

The activation energy and kinetic rates can be further used to interpret the frequency of attempts using the Arrhenius equation:

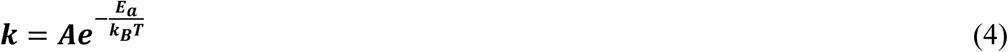

in which the frequency factor *A* corresponds to the frequency of attempts to cross the channel, *k* is the translocation rate (as calculated in Figure 4F), and *E*_*a*_ is the activation energy (as calculated in Figure 4G).

The frequency factors for transitions from pore 1 to 2 (*A*_1_) and from 2 to 1 (*A*_2_) are identical at 1.17 s^-1^. Given the 1 kHz sampling rate of the trajectories, this can be translated to 0.117 % of trajectory localizations within each confinement that can be considered attempts at crossing the narrow channel. This attempt frequency can be translated to a distance from the opening of the channel. To determine the distance from the channel that a particle must be to attempt a hop, we examined the cumulative probability of observing a trajectory point as a function of distance from the channel (**Figure 5**A). It can be found that the distance at which the cumulative probability reaches 0.117% occurs at 15 nm from the channel for pore 1, and 18 nm from the channel for pore 2, noticeably close to the step size of diffusion within the confinement (13 ^[8, 18]^ nm, median [Q1, Q3]). The difference between these attempt distances may reflect different channel geometries on either side.

**Figure 5.**
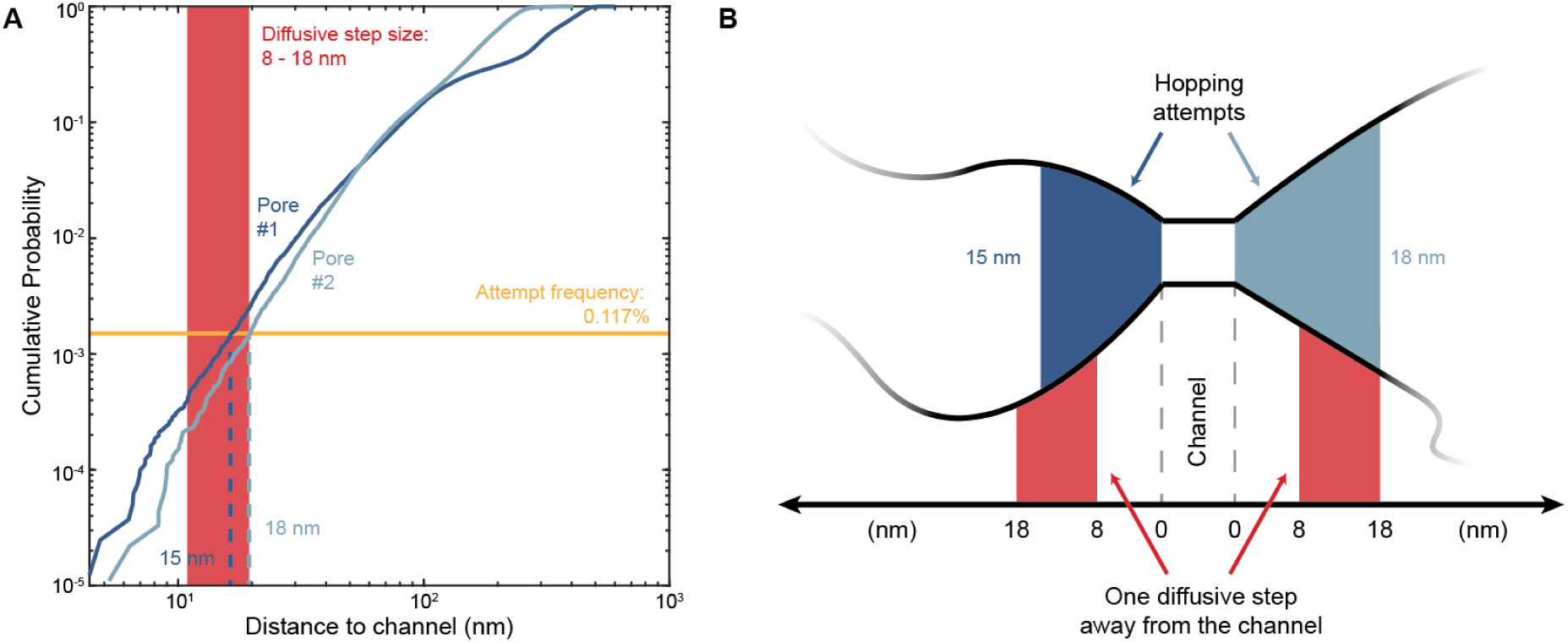
(A) Cumulative distribution function (CDF) of particle localizations as a function of distance to the channel for pore 1 and 2. The solid yellow line indicates the 0.117% cumulative probability corresponding to the frequency factor of 1.17 s^-1^ with 1 kHz sampling. The corresponding distance for hopping attempts is 15 nm and 18 nm for confinement 1 and 2, respectively, as noted by the blue lines. (B) Illustration showing hopping starts to attempt roughly one diffusive step away from the channel.

The measurement of free energy differences and energy barriers can be applied to more complex hopping diffusion scenarios involving multiple connected cavities, by identifying the interconnected cavities and analyzing particle occupation. The trajectory shown in Figure 3A-B was used as an example, and the resulting energy landscape is presented in **Figure 6**A. Building upon this example, we applied the same measurement approach to a larger dataset of 200 nm probes diffusing within 2.0 wt% agarose gel (characterized from 78 hop events across 18 trajectories of 93 total, Figure 6B).

**Figure 6.**
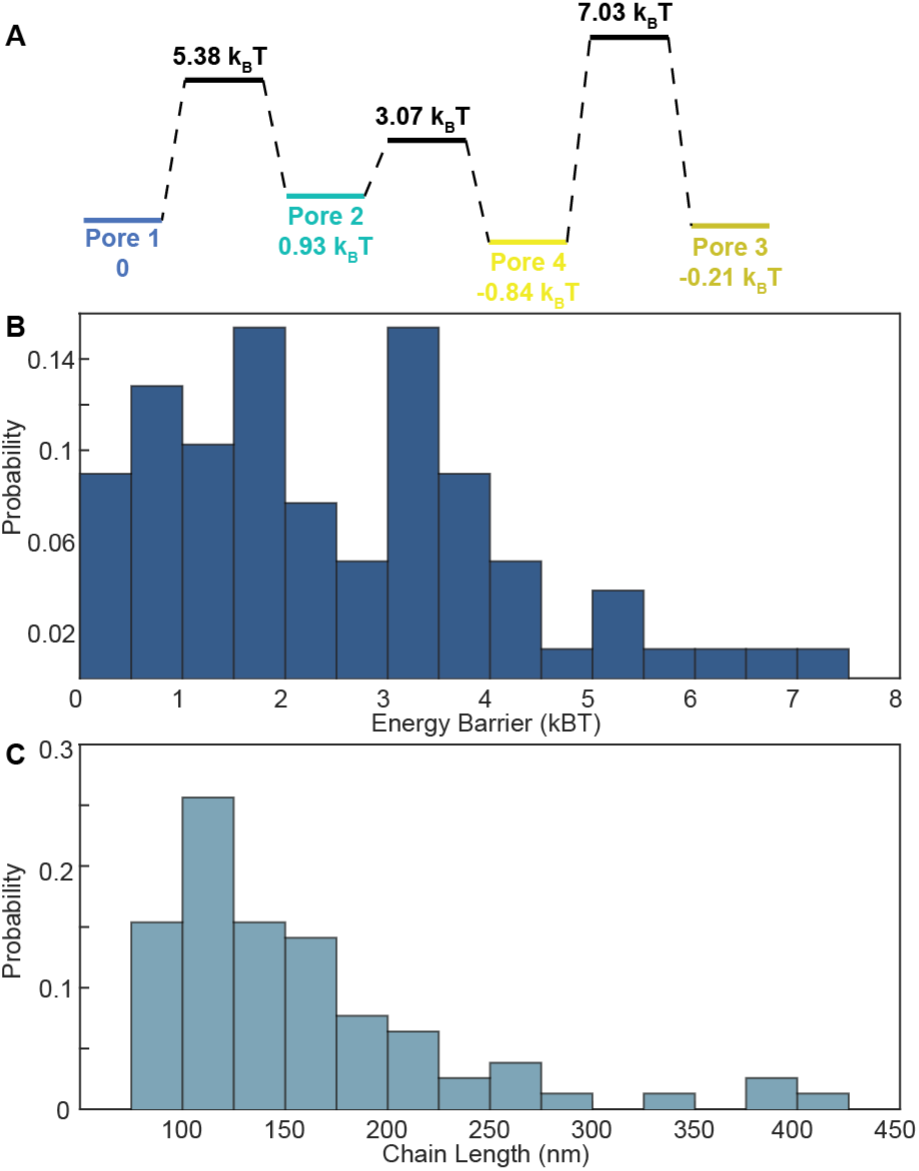
(A) Example of free energy landscape measurement of hopping diffusion across multiple connected cavities, using the trajectory shown in Figure 3A-B. The font color is correspond to cavities identified in Figure 3. (B) Distribution of hopping diffusion free energy barrier of 200 nm probes in 2.0 wt% AG. 78 hops were characterized across 18 trajectories that included hopping events (of 93 trajectories total). (C) Distribution of of chain length calculated via Eq. 5 from the free energy barriers shown in (B).

The free energy barrier involved in hopping diffusion can be explained by different frameworks, including the concept presented in “hopping diffusion”^[51–53]^ and “diffusive escape”.^[54]^ Prior work on hopping diffusion within gel networks describes the fluctuation of a gate between two neighboring confinements, which could allow a particle with size exceeding the network mesh size to hop into a neighboring confinement. In this case, the channel represents the polymer chain fluctuation of the gate, while its volume represents the number of microstates. Based on this hopping diffusion model, the activation energy barrier is given by

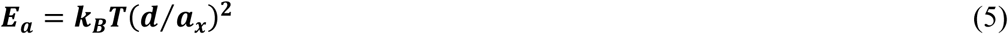

where *d* is the particle size, and *a*_*x*_ is the length of the polymer network.

Based on this equation, the hopping energy barrier of 200 nm probes within 2.0 wt% AG (Figure 6B) can be used to probe the length of polymers within the network. The energy barriers calculated based on particle occupation of the channel results in a calculated network length scale of 130 [107, 181] nm (Median [Q1, Q3], Figure 6C), which is consistent with expected length of 50 nm to >200 nm for 120 kDa agarose.^[55, 56]^ While agarose is a highly heterogenous system, the experiments presented here are a good starting point to understand hopping diffusion in more controlled polymer networks and synthetic films with known (or unknown) structure.

“Diffusive escape” describes Brownian particle escape from confinement through a nanochannel and is usually probed with specially synthesized films, such as a liquid-filled inverse opal film.^[54, 57]^ In the diffusive escape model, the narrow channel itself effectively acts as a free energy barrier, owing to its limited size, as well as additional repulsive potentials that may prevent translocation through the channel. From the current data, if one assumes that the channel exists physically and is rigid, it is possible to estimate the channel volume based on the localization within the channel by *V* ≃ 2std(*x*) × 2std(*y*) × 2std(*z*) (±1 σ, containing ∼ 68% of the data), which gives a result of 5.28 ± 1.24 × 10^5^ nm^3^. Plugging that the volume size of pore 1 (0.012 ± 0.0010 μm^3^ calculated above) into Equation 3 yields an estimated activation barrier of 3.20 ± 0.24 *k_B_T*, which is significantly lower than the value of 5.05 *k_B_T* obtained above from localizations within the channel (Figure 4G). The discrepancy between these two values indicates that an additional energy barrier exists, in addition to the entropic barrier from the limited channel size. This suggests the presence of additional repulsions that hinder translocation beyond the effect of geometry. Notably, this extra barrier appears to be specific to narrow channels, as larger pores exhibit energy differences dependent solely on their volume. This additional obstacle could stem from the hopping diffusion model discussed above, where polymer chain rearrangement introduces an additional energy barrier. Another possible factor is the electrostatic repulsion between negatively charged particles and the negatively charged gel microenvironment, which would be most pronounced at the interface between channel and pores.

More generally within this model, additional free energy barriers in the channel can be categorized into two types based on their effect on particle dwell time. The first type arises from repulsive interactions, such as polymer stretching or electrostatic repulsion, which increase the effective free energy barrier either by introducing steric or energetic obstacles that facilitate faster passage through narrow channels. The second type could theoretically stem from attractive interactions, such as adhesion between the particle and the surrounding polymer networks, which increases the particle dwell time and effectively lowers the net energy barrier imposed by volumetric confinement.

In practice, however, strong adhesion could result in frequent particle immobilization, complicating the entropic landscape with additional microstates and deviating from our current model. Future work will extend this framework to non-agarose systems with controllable interior architectures, such as synthetic systems with tunable pore size, charge distribution, or polymer flexibility. Such studies, especially using particles with varied surface chemistries, would help elucidate how electrostatic or specific interactions modulate hopping energetics.

## 3. Conclusion and discussion

In this work, we have demonstrated that active-feedback single-particle tracking methods have great potential for microrheology studies, providing highly spatiotemporally resolved 3D information. The combined high sampling rate, 3D localization, and long duration of observation allowed for the complete characterization of nanoscale confinements and channels within heterogeneous agarose gel networks. Although hopping diffusion has been previously reported, this work, for the first time, quantitatively characterizes the duration of hopping events and extracts free energy barriers directly from single-particle trajectories. The direct observation of these hopping events between two adjacent nanoscale pores presents a unique opportunity to understand narrow escape events in hydrogel networks. Furthermore, the high spatiotemporal resolution of each single-particle trajectory also serves as structural probes, mapping the interior microstructure of 3D porous materials in detail (Figure 3 and Figure 4E).

In this work, we also presented a novel analytical method to resolve nanoscale structures from 3D trajectories, which is necessary to understand diffusion heterogeneity originating from structural heterogeneity. In this analytical method, neighboring localizations are grouped together to interpret structural information from highly dynamic diffusing but ergodic trajectories. Sufficient localization sampling and high temporal resolution is a critical prerequisite for such an analytical method, which is, to our knowledge, only fulfillable by active-feedback methods such as 3D-SMART.

This work also demonstrates the broader potential of active-feedback tracking for probing transport and structure in a wide range of soft and heterogeneous materials. Placeholder for #2-4 This technique will also present new avenues to study the penetration of virus or drug delivery cargo through mucus or mucin hydrogels,^[38, 58]^ interpreting the microscopic view of particle dynamics during gel electrophoresis or column chromatography,^[14, 59]^ exploring the mechanism behind critical material chemistry, such as self-healing and catalytic processes,^[60, 61]^ and illustrating the dynamics within biomolecular condensates.^[62]^

## 4. Methods

### Sample preparation

Agarose powder (type I low EEO, Sigma-Aldrich) was mixed with 1× Tris-Acetate-EDTA buffer (Apex Bioresearch Products, 18-134L) in a microwaveable flask and microwaved until the agarose was completely dissolved without overboiling. Cool down the agarose solution down to about 60 ℃ in water bath. 100 nm, 200 nm, or 500 nm carboxyl-functionalized polystyrene microspheres fluorescently labeled with Dragon Green (Bangs Laboratories, Inc., FCDG002, FCDG003, or FCDG005) were pre-diluted in distilled water and pre-warmed to about 60 ℃ in water bath. The anionic carboxyl group on the emitter beads would be expected to have minimal interaction with the anionic agarose.^[63]^ Also, it was demonstrated that premixing probes with agarose gel has no significant impact in gel formation.^[39]^ Equal volume of agarose solution and microsphere dilution were quickly mixed to achieve the desired final concentration. 300 μL of mixture was transferred into a chambered coverslip (Ibidi, 80826). The sample was solidified under room temperature for about 15 minutes and 100 μL distilled water was applied on the top of gel to prevent dehydration.

### Validating the tracking of an individual particle

As part of the 3D-SMART mechanism, the EOD continuously scans the particle in a 5×5 grid. The result is a real-time “EOD image” which can be used to evaluate tracking performance. If only one individual particle is being tracked, the center pixel of the EOD image will have most photon counts. If more than one particle exists in the scanning plane, multiple photon-count maxima will exist in a single EOD image, which is also accompanied by an increase in the average particle intensity. These two factors can be used to rule out trajectories containing multiple particles. The intensity recorded during the trajectory can be used to confirm a single particle is being tracked, and to further identify when he system “jumps” from one particle to another. The stability of the intensity is evaluated by the ratio of the standard deviation and mean of the intensity in each 1-sec window. A ratio larger than 0.2 (intensity 50% larger or 33% smaller than the mean) indicates a new particle entering the observation volume. These transitions are used to break trajectories into segments containing only one particle. An example of the system tracking more than one particle is shown in the Supporting Information (Figure S3).

### Cavity identification

For 2D cavity identification, trajectories were processed using a sliding window of 180 second window, stepped in 30-second increments. Particle localizations were distributed into 10×10 nm^2^ pixels, selected based on the localization precision. The density of localizations in each pixel is convoluted with a 3×3 Gaussian Kernel. Pixels with fewer than 15 localizations were filtered out. Areas were grouped using the `bwconncomp` function in MATLAB. Areas in different segments were merged to form the final result. Areas with fewer than 60 pixels were excluded. This threshold was determined by observing fixed particles.

Similarly, in 3D cavity identification, trajectories were processed using a sliding window algorithm, with 180-second window size and 30-second step size. Particle localizations were distributed into voxels with the size of 10×10×20 nm in XYZ, determined by localization precision. The localization density in each voxel is convoluted with a 3×3×3 Gaussian Kernel. Voxels with fewer than 2 localizations were filtered out. Volumes were grouped using the `bwconncomp` function in MATLAB. Volumes in different segments were merged to form the final result. Final volumes with fewer than 300 voxels were excluded. This lower threshold was determined from trajectories of fixed particles. The volumetric images were plotted with the `vol3d` function.^[64]^

The minimal pixel/voxel occupancy (15 for 2D and 2 for 3D) mentioned above was determined by applying the algorithm to 3D simulated trajectories, with duration of 180 seconds, time steps of 1 millisecond and diffusion coefficient of 0.03 μm2/sec (the diffusion coefficient of 200 nm probes in 2 wt% AG), and 10 nm (XY) and 30 nm (Z) localization precision in XY and Z respectively to mimic slower but unconfined diffusion. In this case, identified pixels represent apparent “confinement” from random motion. The cumulative density functions of the number of dwells in 2D pixels and 3D voxels are shown in Figure S15. For 2D analysis, only 5% of pixels have pixel occupancy greater than 15; therefore, this value was chosen as the threshold for defining confined diffusion. For 3D analysis, a minimum pixel occupancy of 2 represents 92% of all simulated localizations, so this was set as the threshold for determining a legitimate dwell.

The pixel/voxel number within each isolated area/volume represents the cavity area/volume size, while numbers of particle localizations within the area/volume represent the dwell time within the area/volume. Measuring uncertainty of the area/volume size is calculated by expanding or shrinking the area/volume by one pixel/voxel.

### Position categorization

Positions were categorized based on their spatial relationship to cavities. Positions labeled as ‘not-in-cavity’ were further classified as follows:

#### Successful hop

Consecutive positions labeled ‘not-in-cavity’ that connect two distinct cavities were registered as ‘successful hop’, indicating pathways or connections between cavities.

#### Unsuccessful hop

The rest of the consecutive positions labelled ‘not-in-cavity’ and located within the channel were registered as ‘unsuccessful hop’, indicating events of particle entering the channel but not successfully pass through.

#### Within the channel

Localizations classified as both “successful” and “unsuccessful” hops, as detailed above, are combined to yield the total number of particle localizations within the channel. For example, in Figure S9a, points colored in dark blue and light blue represent the particle localization within two different cavities. The points between the two cavities (colored in red) are temporally resolved into two subcategories: (1) successful hops: points successfully connected two cavities; (2) unsuccessful hops: points entered the channel volume but failed to pass through.

#### Measuring uncertainty

All remaining ‘not-in-cavity’ positions that do not connect cavities are near cavities but lack sufficient sampling to be definitively assigned to any specific cavity. These positions represent ‘measuring uncertainty’. For both thermodynamic and kinetic analysis, these uncertain positions were ignored.

### Fractional dwell time equals fractional confinement volume

For diffusing particle constituting isolated states with total volume of *V_tot_* and total dwell time of *t_tot_*. For each individual state with volume of *V*_*k*_ and dwell time of *t*_*k*_, according to the partition function, the fractional dwell time is

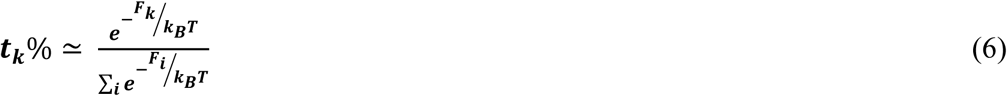

in which, *F*_*k*_ the thermodynamic potential energy. In this scenario, the thermodynamic potential is simply contributed by entropy, *F* = −*ST*, which can be determined using Boltzmann’s entropy formula:

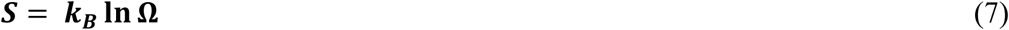

in which, *k*_*B*_ is Boltzmann’s constant, and Ω is the number of microstates within macrostate. Therefore, the fractional time can be written as

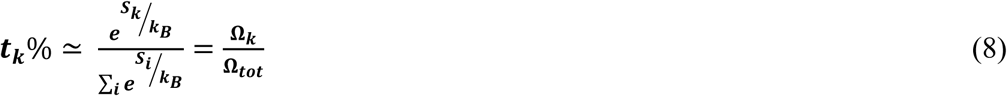

For the isolated confinements constituted by diffusing particles with negligible particle-confinement-interaction, Ω is proportional to the volume of confinements, *V*. Naturally, we have the fractional dwell time equal to the fractional confinement volume.

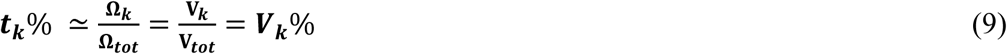

where ***t***_***k***_ is the dwell time and ***V***_***k***_ is the confinement volume in confinement *k*. ***t****_tot_* is sum of all dwell times and *V_tot_* is the sum of all confinement volumes in confinements across each individual SPT trajectories. ***t***_***k***_% is the fractional dwell time and ***V***_***k***_% in each confinement.

## Acknowledgements

We acknowledge financial support from the National Institute of General Medical Sciences of the National Institutes of Health under award number R35GM124868 (K.D.W.). We appreciate the support from the Fitzpatrick Institute for Photonic at Duke University, along with the John T. Chambers Scholarship (to Y.L.). We thank Dr. Michael Rubinstein at Duke University and Dr. Dario Conca at Umeå University for useful discussions. Some schematics were created with BioRender.com.

## Data Availability Statement

Data available from the corresponding author upon reasonable request

Received: ((will be filled in by the editorial staff))

Revised: ((will be filled in by the editorial staff))

Published online: ((will be filled in by the editorial staff))

## Supporting Information

Supporting Information is available from the Wiley Online Library or from the author.

## The table of contents entry

High-speed 3D tracking microscopy enables real-time tracking of nanoparticles in porous hydrogels with high spatiotemporal resolution, revealing diffusion heterogeneity and hopping diffusion events. The quantitative analysis of trajectories uncovers nanoscale pore structures and energy landscapes, providing new insights into nanoparticle transport mechanisms and porous materials at the single-particle level.

## Supporting Information

## 1 Experimental Set-up

### 1.1 Instrumental Set-up Overview

Detailed instrumental diagram was depicted in Figure S1. The tracking emission is merged from the objective and is reflected to the detection pass with a multiband entrance dichroic (**DCM1**, Chroma, no. ZT405/488/635rpc). The beam was then split by a 640 nm short-pass dichroic mirror (**DCM2**, Semrock, no. FF01-640/14-25). The emission filter for the tracking signal detection is a 535/50 bandpass filter (**f2**, Semrock, no. FF01-535/50-25).

### 1.2 3D-SMART data acquisition workflow

The data acquisition workflow is automated. Briefly, it is initiated by searching for a particle using the piezoelectric stage. When the number of detected photons exceeds a predetermined threshold, a particle is assumed to be located within the detection volume, and the active-feedback tracking loop will be triggered to ‘lock on’ the diffusing particle. When the trajectory reaches a set maximum (usually 3 or 5 minutes), tracking will be disengaged, which follows with a new round of searching. The maximum experimental duration was 3 hours to avoid aging and deformation of AG.

## 2 Analytical Methods

### 2.1 Mean-square displacement analysis for particle diffusion in hydrogel

Trajectories were quantified using diffusion coefficient (D), power exponent α, and packing coefficient (Pc). A power-law mean-square displacement (MSD) analysis is typically applied for anomalous diffusion.

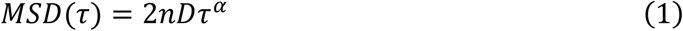

Where n is the number of dimensions, D is the diffusion coefficient, τ is the lag time, and the exponent α is used to classify anomalous diffusion (α < 1: subdiffusion; α = 1: Brownian motion; α > 1: superdiffusion).

However, in practice, MSD has limitations when exploring highly confined trajectories. While α can be used to characterize sub-diffusion, experimental values for α exhibit a wide and noisy distribution for highly diffusive particles. On the other extreme, for groups with lower diffusivities, α varies little and is not very informative (Figure S6c&f).

### 2.2 Packing coefficient analysis for particle diffusion in hydrogel

An alternative method to quantify lateral subdiffusion is the packing coefficient (*Pc*), which was demonstrated by Triller et al. to be a powerful tool in distinguishing confinements. It compares the lateral displacements in a time window and the occupied surface area:

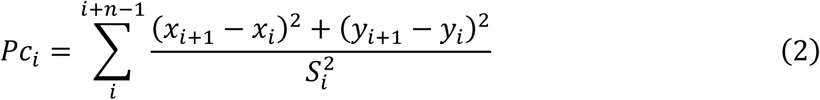

where x_i_, y_i_ are the coordinates at time i; n is the length of the time window; and S_i_ is the surface area of the convex hull of the trajectory segment between time points i and i+n. A larger packing coefficient means a higher degree of confinement. A window of n = 1000 time points (1 second) was used throughout the analysis unless otherwise indicated.

Packing coefficient analysis showed a clear increasing trend of confinement with increasing concentration of AG and probe size (Figure S6a&d), in agreement with prior reports that higher agarose concentrations result in smaller pore sizes in the gel.

### 2.3 Pore size calculations using packing coefficient

One useful feature of Pc is that it only scales inversely to the size of the confinement area A and independently of D, providing a measure of confinement degree in environments with different viscosities and a different method to calculate the size of confinements. Trajectories of particle diffusing within confinements of certain sizes were simulated, and the relationship between Pc and A was empirically derived as:

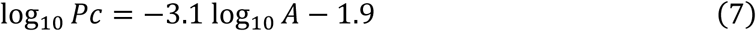

The 2D area size can be calculated accordingly. Figure S11 shows a very similar result compared to the calculation from the 2D heatmap in Figure 4E.

### 2.4 Monte Carlo simulations of free or confined diffusion

The program was written in MATLAB (The MathWorks, Natick, MA). The displacement in x and y were generated randomly from a normal distribution with the mean of zero and the variance equal to 2Dτ (τ = 1 ms). In the case of confined diffusion, particle diffusions start from the center of the confinement area. When the particle moves out of the confinement area, a normal vector at the point of collision will be calculated and reflect the displacement vector accordingly. This ‘reflection’ repeats until the particle fails within the confinement area. The localization noise was simulated by adding a distance to displacement in x and y, where the distance is randomly generated from an independent normal distribution with a mean of zero and a given variance corresponding to the localization precision obtained experimentally.

### 2.5 Additional hopping example of 500 nm probe within 2.0 wt% AG

The occurrence of hop events is strongly related to the mesh size and probe size, as demonstrated by Cai et al. Due to the wide distribution of mesh sizes in agarose gel, hopping diffusion could also occasionally happen with 500 nm probe in 2 wt% AG (2 out of 57 trajectories, Figure S12). Due to the combination of the larger probes and relatively dense meshes, the relocation rate is much slower than that previously shown for 200 nm probes.

### 2.6 Evaluating the tracking performance during hops

To further confirm the directionality of hops and exclude artifacts that the lag of the piezoelectric stage may cause, we evaluated the tracking performance by comparing the central photon fraction. In short, we used a short bin time for the EOD photon arrival image, 0.5 ms, and for each image, the central photon fraction was calculated as the central photon counts divided by the sum of all photons.

We compared photon arrival information during hops with 5-sec random non-hop sections from the same trajectory. The distribution of central photon fraction shows no significant difference (p = 0.56), which indicates tracking performance remained the same during hops.

## 3 Figures and Movies

**Figure S1:**
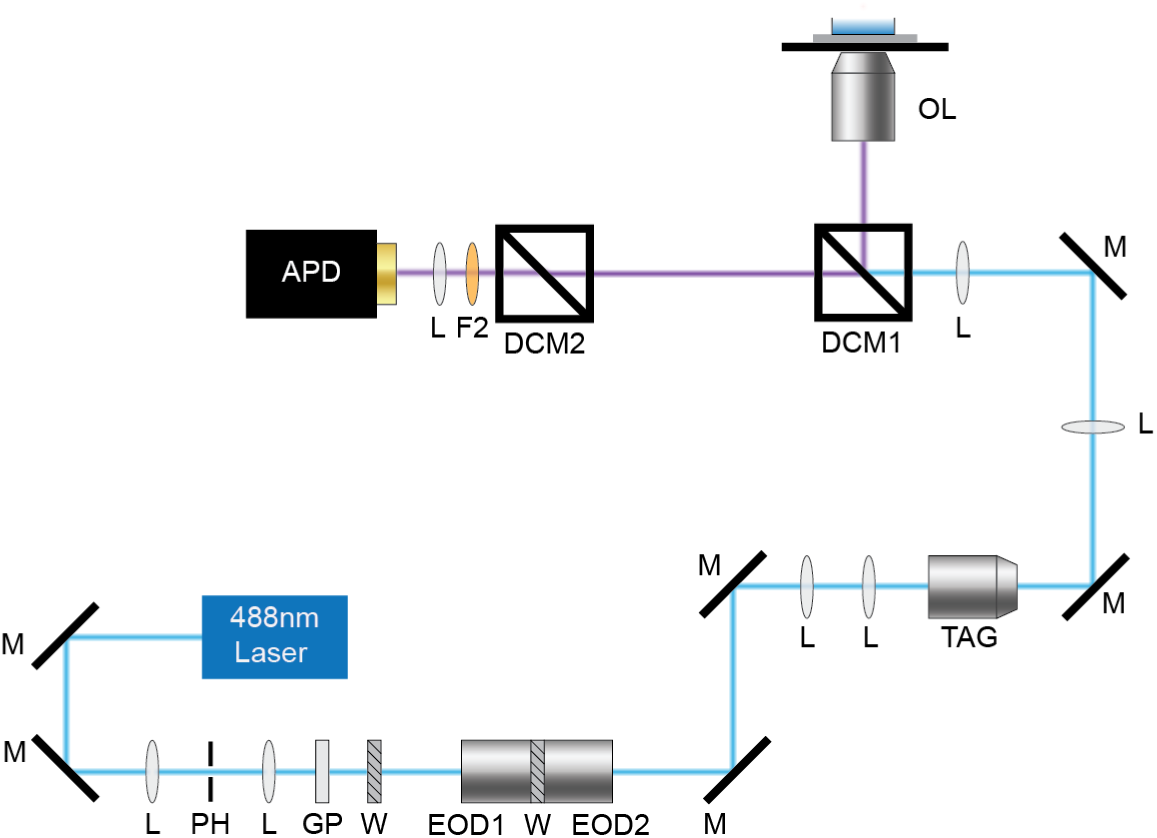
Instrument diagram. M: mirror, L: lens, PH: pinhole, GP: Glan-Thompson Polarizer, W: half wave plate, EOD: electro-optic deflector, TAG: tunable acoustic gradient lens, DCM: dichromatic mirror, OL: objective lens, F: fluorescence emission filter.

**Figure S2:**
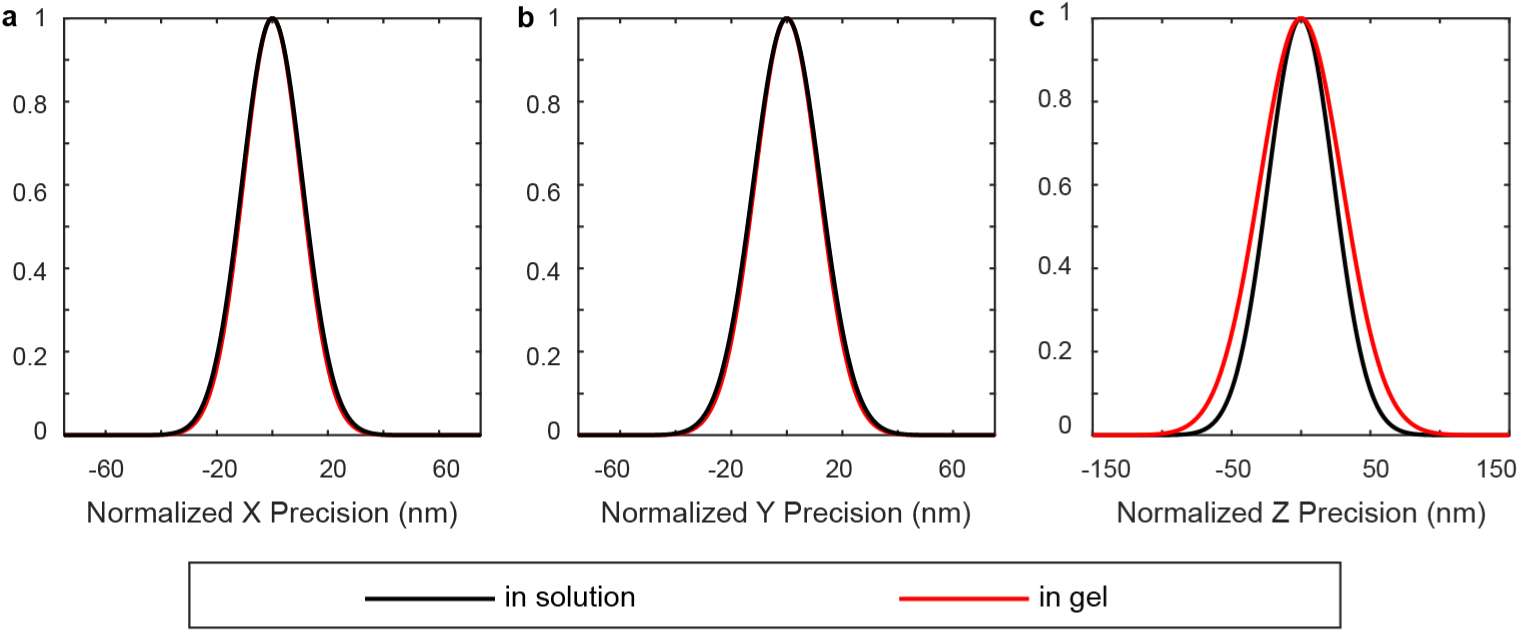
Tracking precisions of 3D-SMART. Tracking precisions were calculated from two trajectories of fixed fluorescent microspheres with similar intensity (∼ 300 kHz). (**a**-**c**) Standard deviations of X, Y and Z position readouts (in 1 sec segments) in solution (black) and in agarose gel (red). The lateral localization precisions in solution and in gel are similar (in solution: X: 10.5 ± 0.6 nm, Y: 11.6 ± 0.4 nm; in gel: X: 10.9 ± 0.5 nm, Y: 12.0 ± 0.4 nm). The axial localization precision in gel (29.7 ± 2.8 nm) is slightly impacted by scattering compared to precision in solution (23.7 ± 1.3 nm).

**Figure S3:**
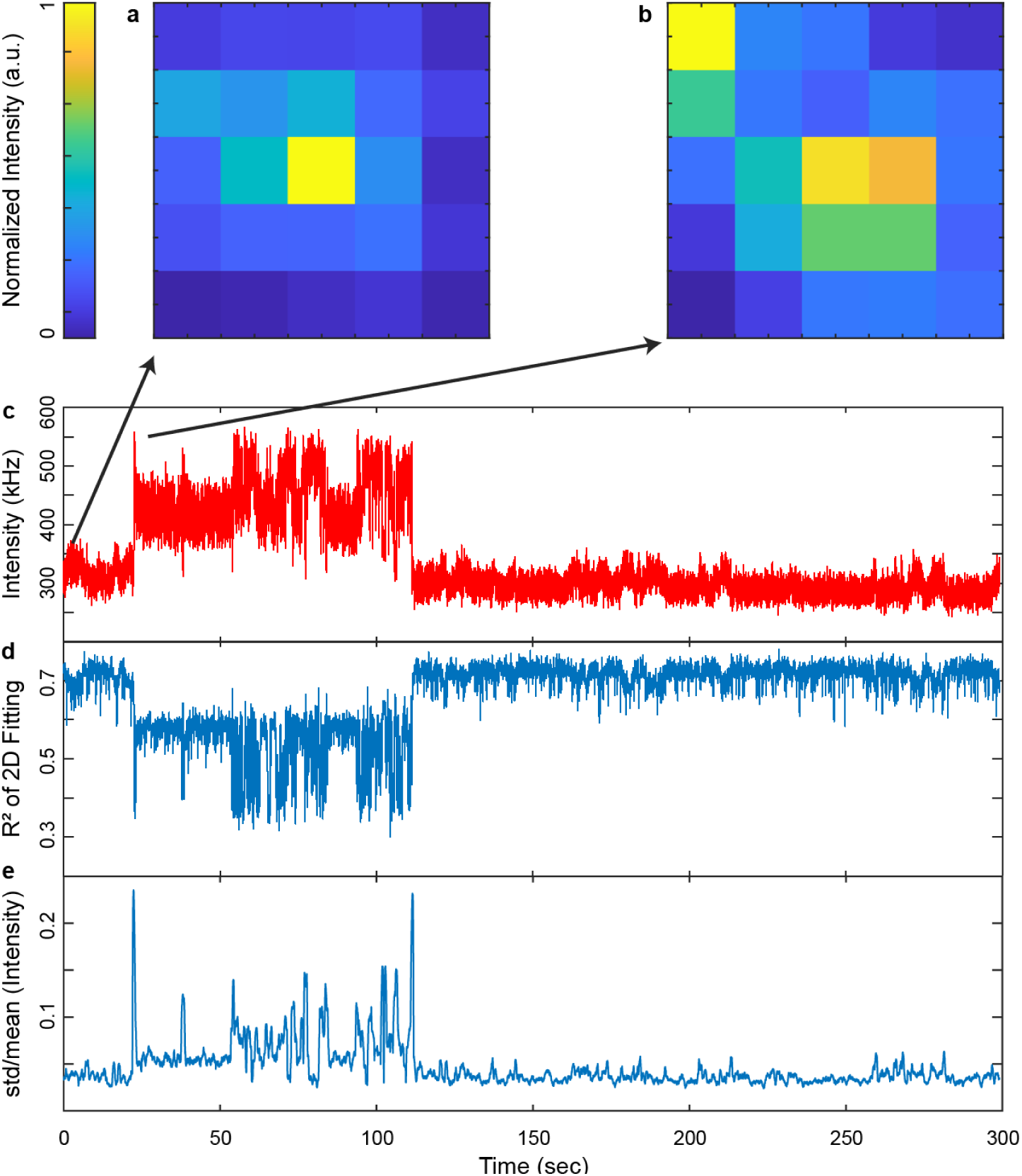
Validation of single-particle tracking. (**a**) At the start of the trajectory a single particle is being tracked as evidenced by the single peak in the observation volume. (**b**) At ∼ 20 seconds, a second particle enters the scanning plane from the left-top corner. (**c**) The arrival of the second particle can be observed in the intensity trace. (**d**) The second particle can also be detected by the goodness of a unimodal 2D Gaussian fit of the detected photon distribution. (**e**) The intensity stability (ratio of standard deviation to the mean of the intensity) can be used to identify the beginning and end of multi-particle trajectory segments.

**Figure S4:**
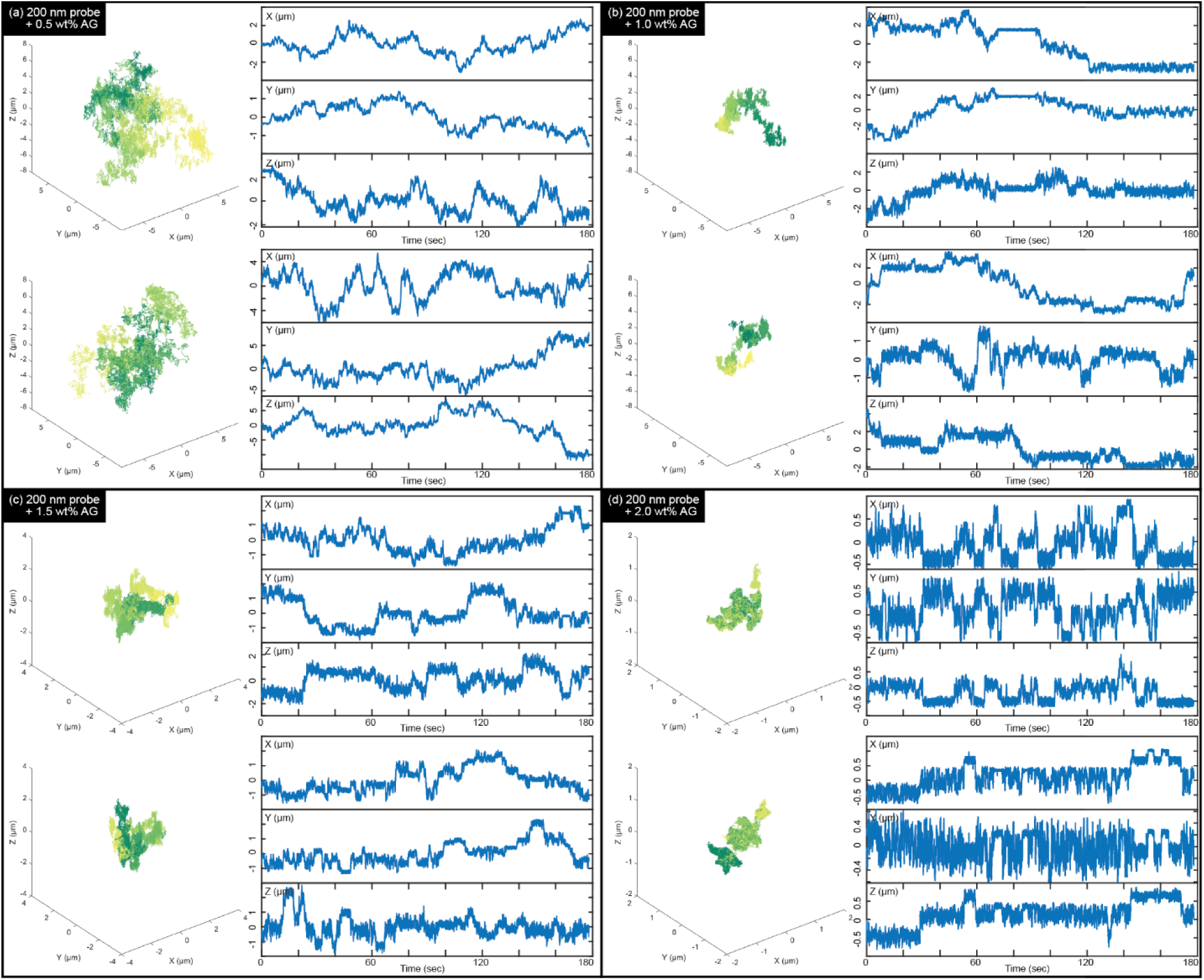
Trajectory, X, Y, and Z trace of 200 nm probes within increasing wt% AG, related to Figure 2A. Trajectory, X, Y, and Z trace of 200 nm probes with (**a**) 0.5 wt% AG, (**b**) 1.0 wt% AG, (**c**) 1.5 wt% AG, and (**d**) 2.0 wt% AG.

**Figure S5:**
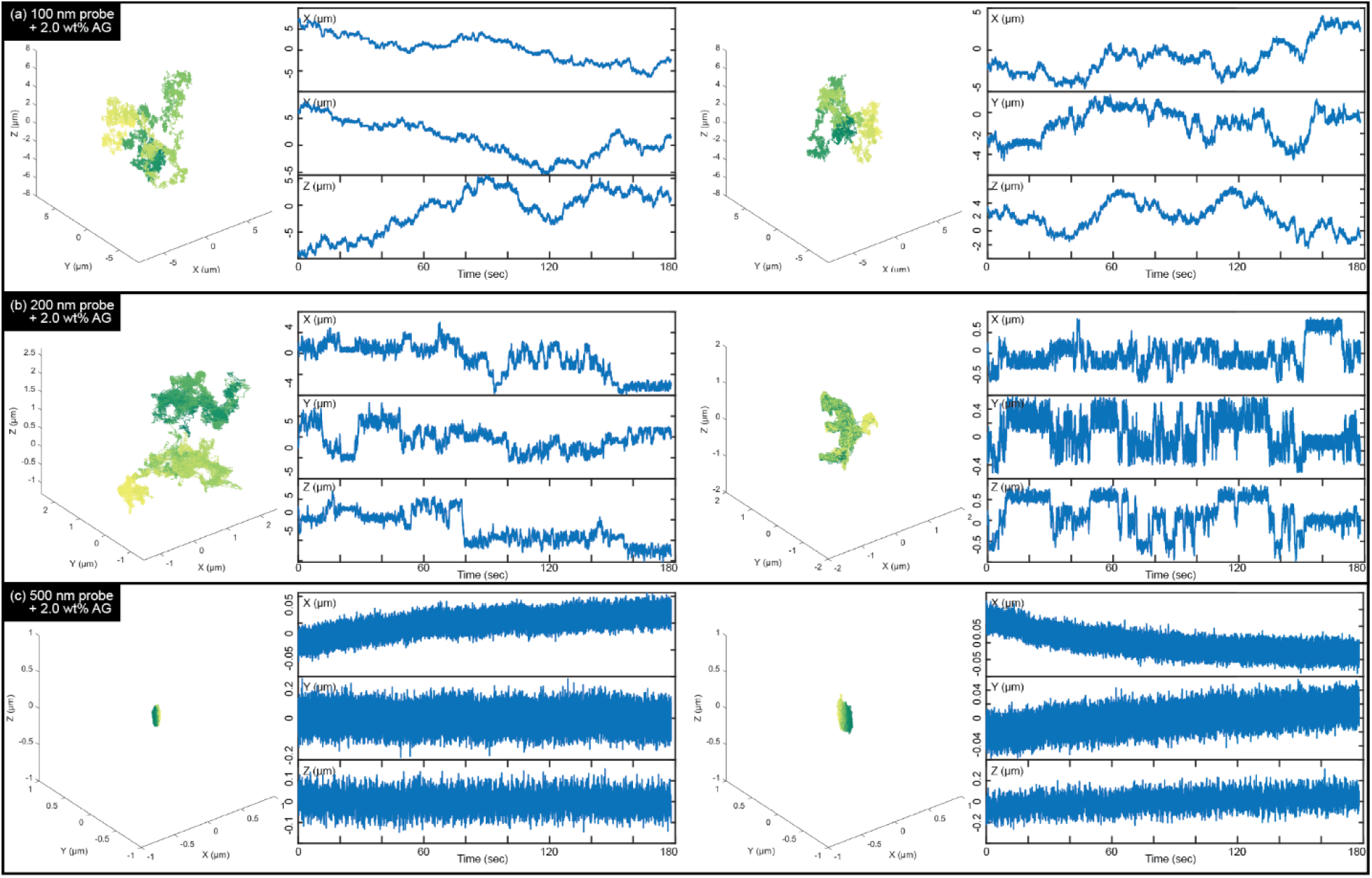
Trajectory, X, Y, and Z trace of different sized probes within 2.0 wt% AG, related to Figure 2B. Trajectory, X, Y, and Z trace of (**a**) 100 nm probes, (**b**) 200 nm probes, and (**c**) 500 nm probes diffusing within 2.0 wt% AG.

**Figure S6:**
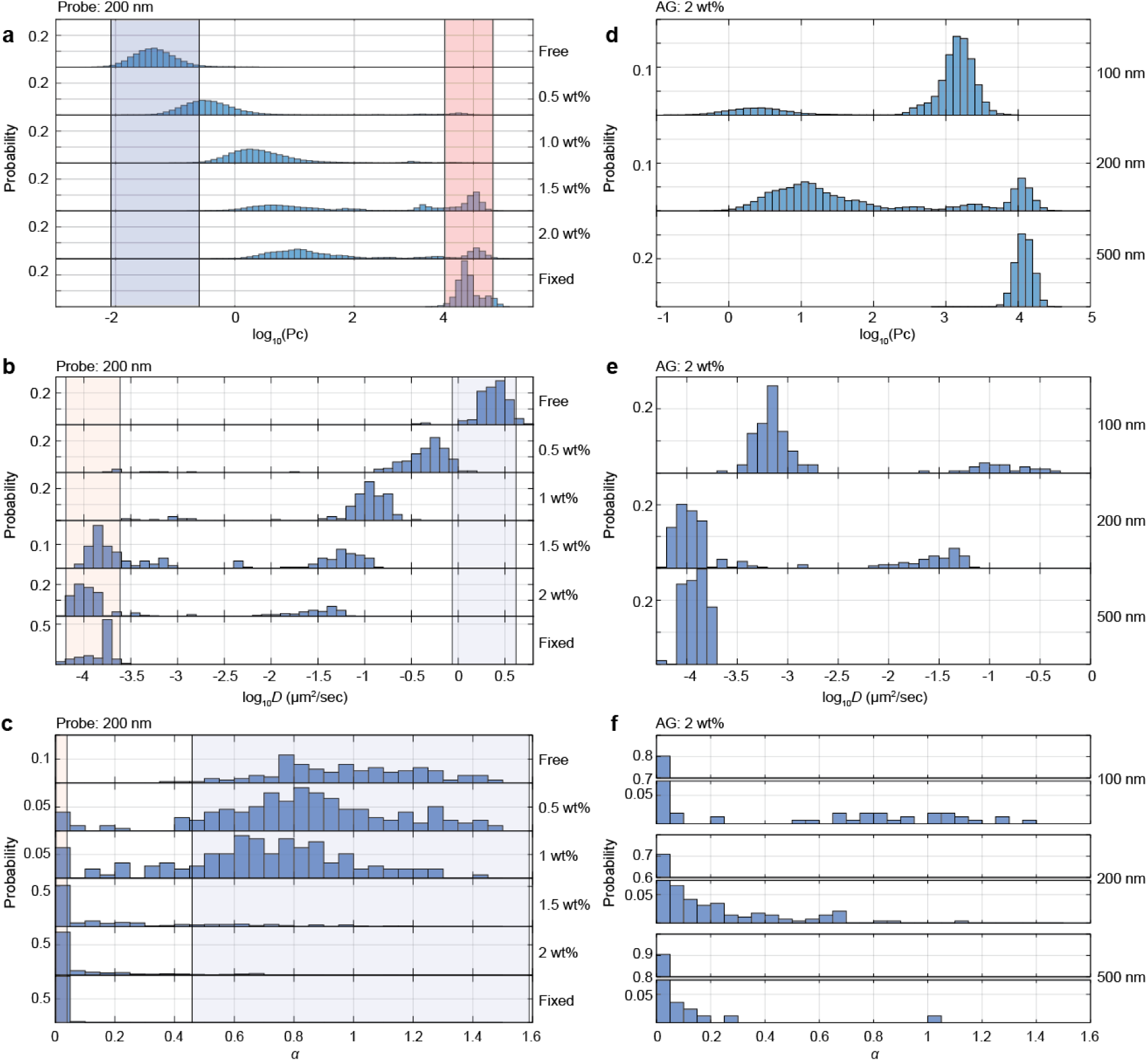
Diffusion coefficients and alpha of trajectories with different probe size and AG concentration, related to Figure 2. (**a**) Distribution of packing coefficients of 200 nm probes diffusing in water, increasing wt% AG, and fixed on the surface of the coverslip, respectively. The range of Pc is highlighted in red for fixed 200 nm probes (mean ± 2×std) and purple for freely diffusing 200 nm probes (mean ± 2×std). (**b**-**c**) Distribution of diffusion coefficients and exponential exponents α of 200 nm probes diffusing in water, increasing wt% AG, and fixed on coverslip, respectively. The range (mean ± 2×std) of diffusion coefficients and α corresponding to fixed trajectories is highlighted in light red and the range of free diffusing trajectories is highlighted in light purple. (**d**) Distribution of packing coefficients of different-sized probes diffusing in 2.0 wt% AG. (**e**-**f**) Distribution of diffusion coefficients and exponential exponents α of 100, 200, and 500 nm probes diffusing in 2.0 wt% AG, respectively.

**Figure S7:**
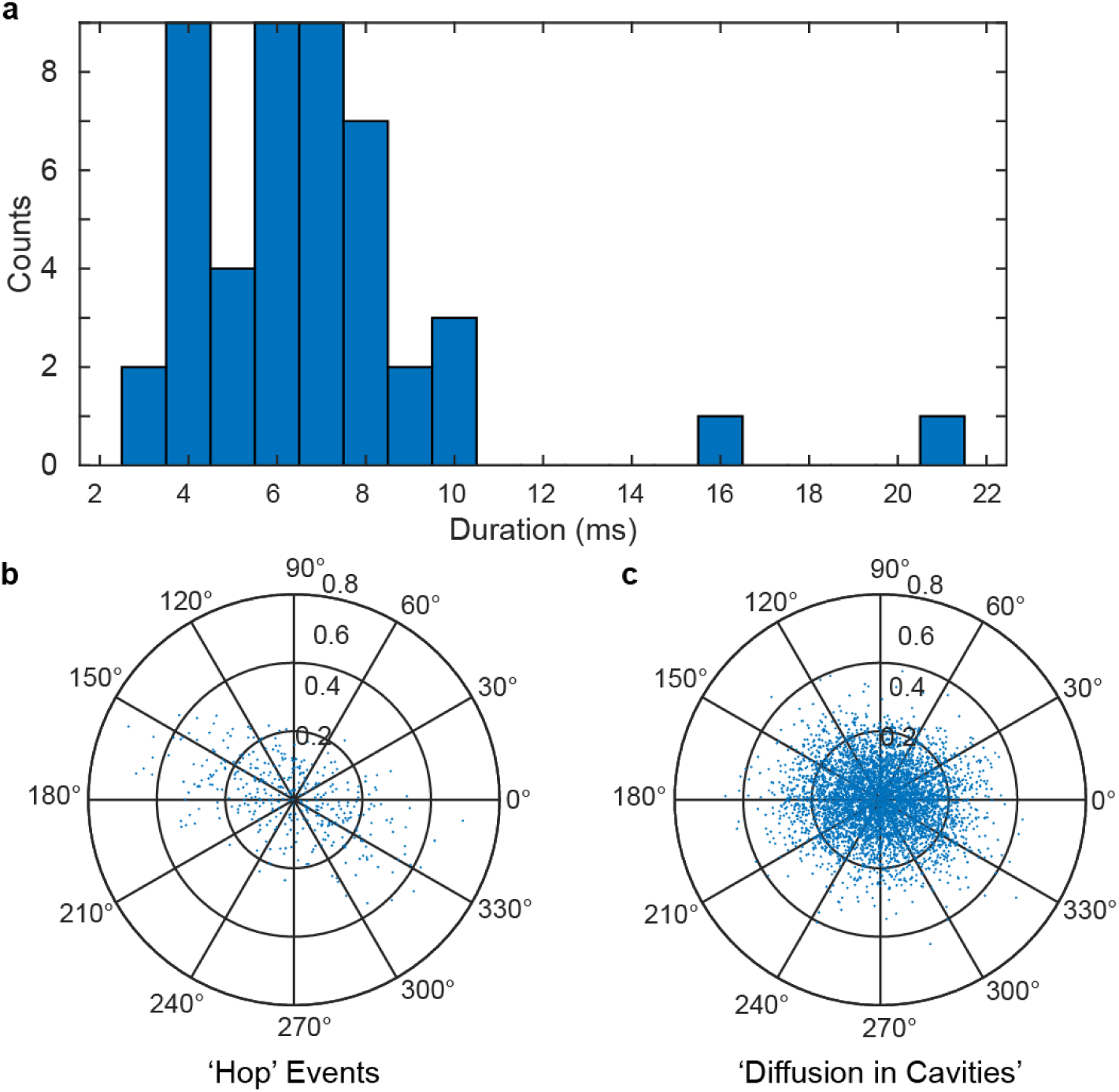
Hop events are momentary, highly one-dimensional, faster, related to Figure 4. (**a**) Typical escape event durations were 6.3 ± 1.9 ms with two outliers of 16 and 21 ms, respectively. These momentary events happen faster than the frame rate of most image-based tracking methods. *N* = 42. (**b** & **c**) Polar scatter plot (direction of displacement and magnitude of displacement) of ‘hop’ events and diffusion within cavities, respectively.

**Figure S8:**
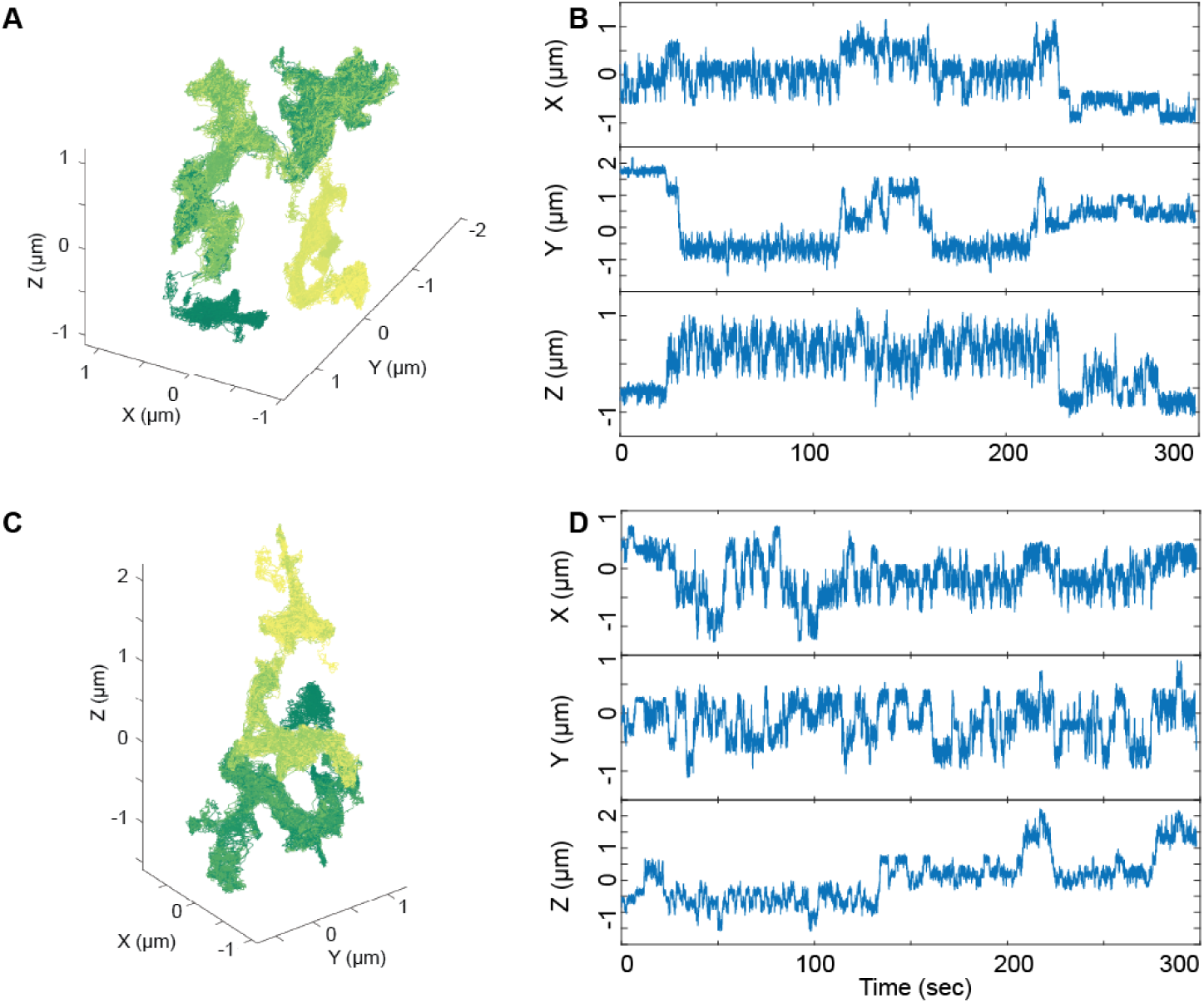
Detailed information of X, Y, and Z trace, related to Figure 2C-E. (**A** and **B**) 3D trajectory, X, Y, and Z trace of the trajectory shown in Figure 2C-D. (**C** and **D**) 3D trajectory, X, Y, and Z trace of the trajectory shown in Figure 2E.

**Figure S9:**
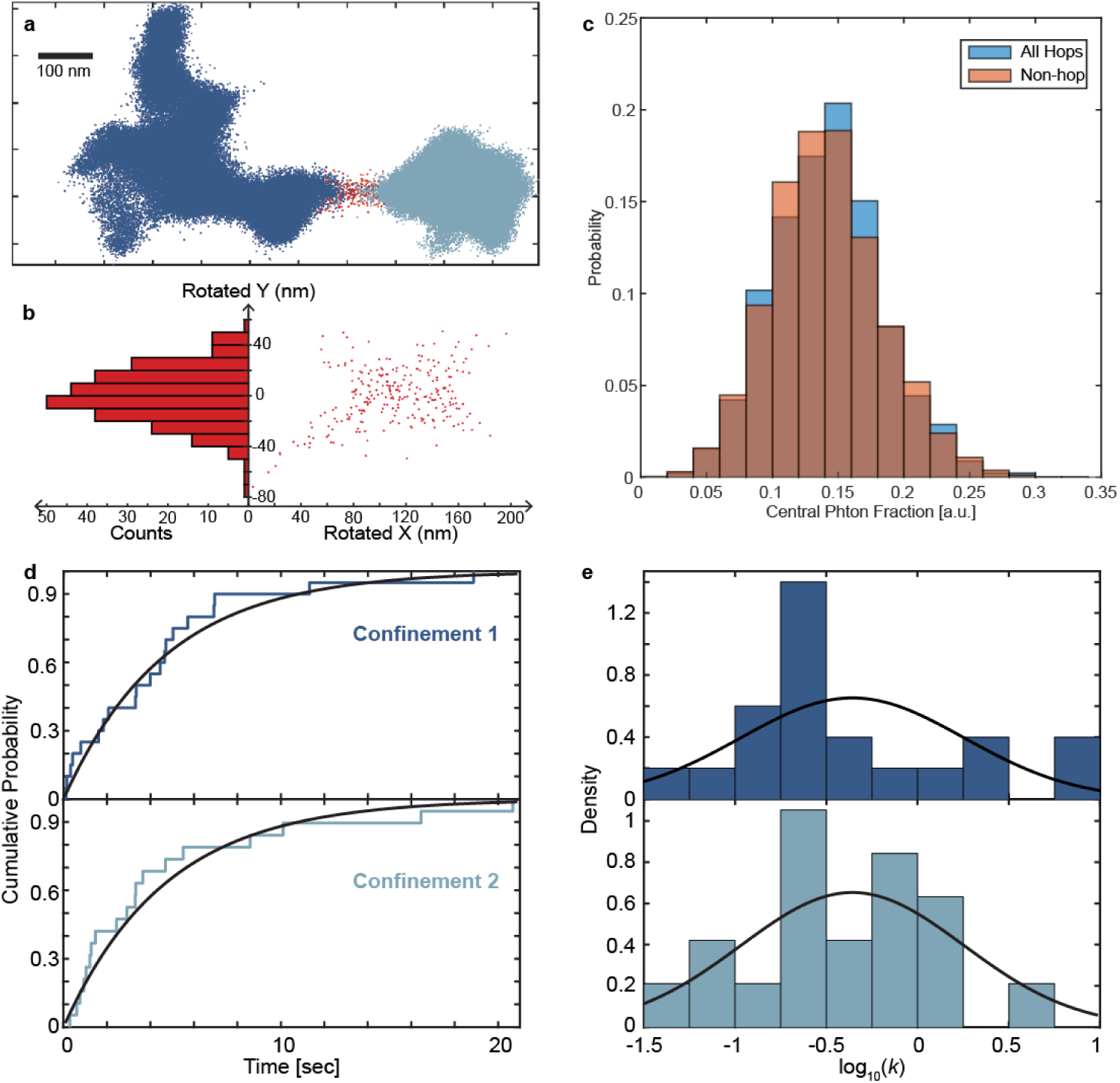
Supplementary figures related to Figure 4. (**a**) Rotated view of point cloud of localizations. Dark blue points and light blue points represent two populations, and red points represent the transition between two states. (**b**) Right: rotated coordinates of the transition points. Left: histogram of the rotated Y position of the transition points as a measurement of the channel width, calculated to be 3 × std = 68.0 ± 3.3 nm (calculated value ± measuring uncertainty). (**c**) Central photon fraction distribution of ‘hop’ events and diffusion within confinements shows no significant difference (*p* = 0.55). The bin time was 0.5 ms. This indicates good tracking performance during hops. (**d**) Cumulative probability function of relocation times of confinement 1 (dark blue) and confinement 2 (light blue). Black lines represent exponential distribution fitting. (**e**) Distribution of relocation rates of ‘hop’ from confinement 1 to 2 (dark blue) and from confinement 2 to 1 (light blue). Black lines represent normal distribution fitting.

**Figure S10:**
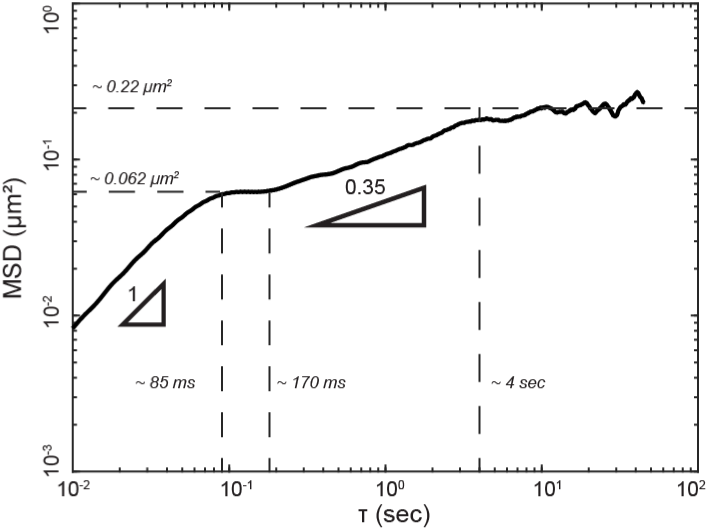
3D MSD plot, related to Figure 4. 3D MSD plot of the representative trajectory shows different diffusive states in cavities connected by a channel (minimum lag time of 10 ms). The two slopes were calculated using a linear fit within the corresponding time window.

**Figure S11:**
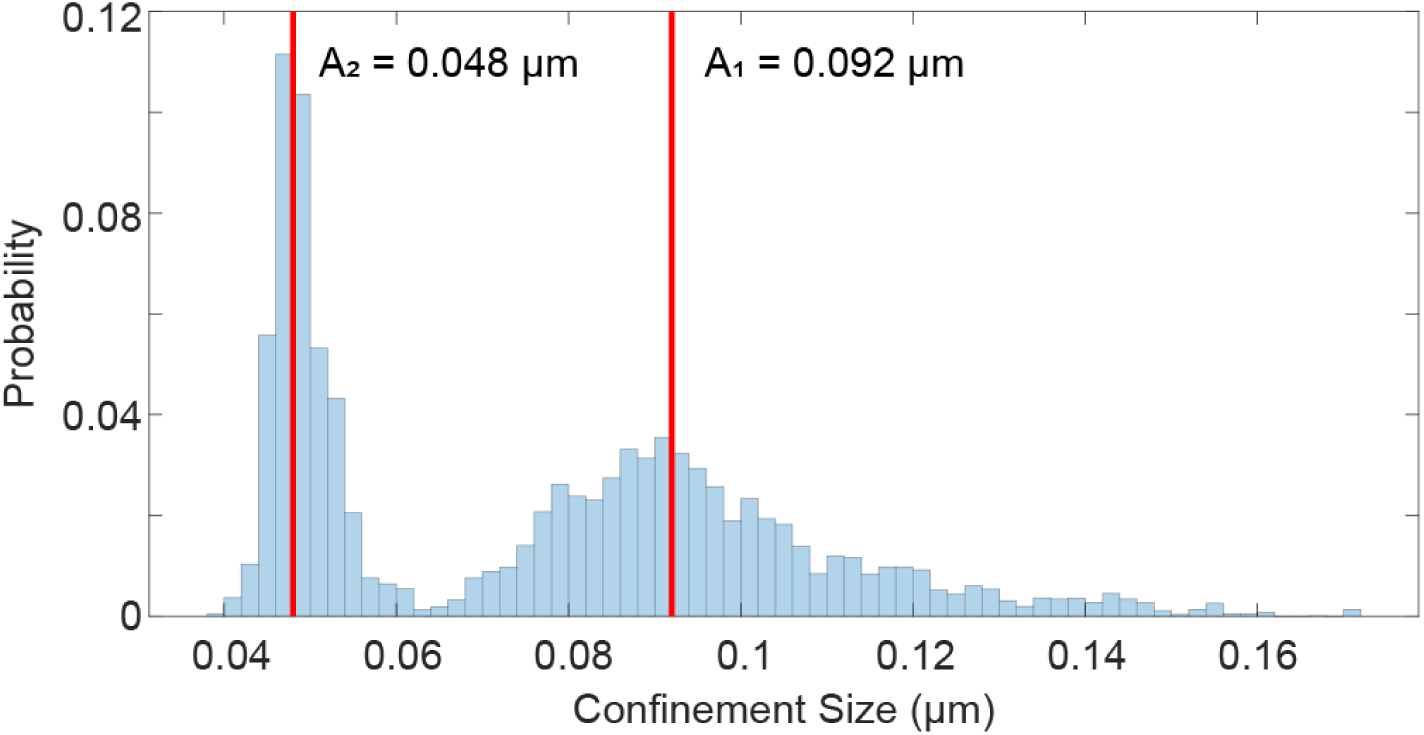
Confinement size calculation through packing coefficient, related to Figure 4. Histogram shows the confinement size calculated from the packing coefficient. The red line shows the confinement size calculated from the 2D heatmap.

**Figure S12:**
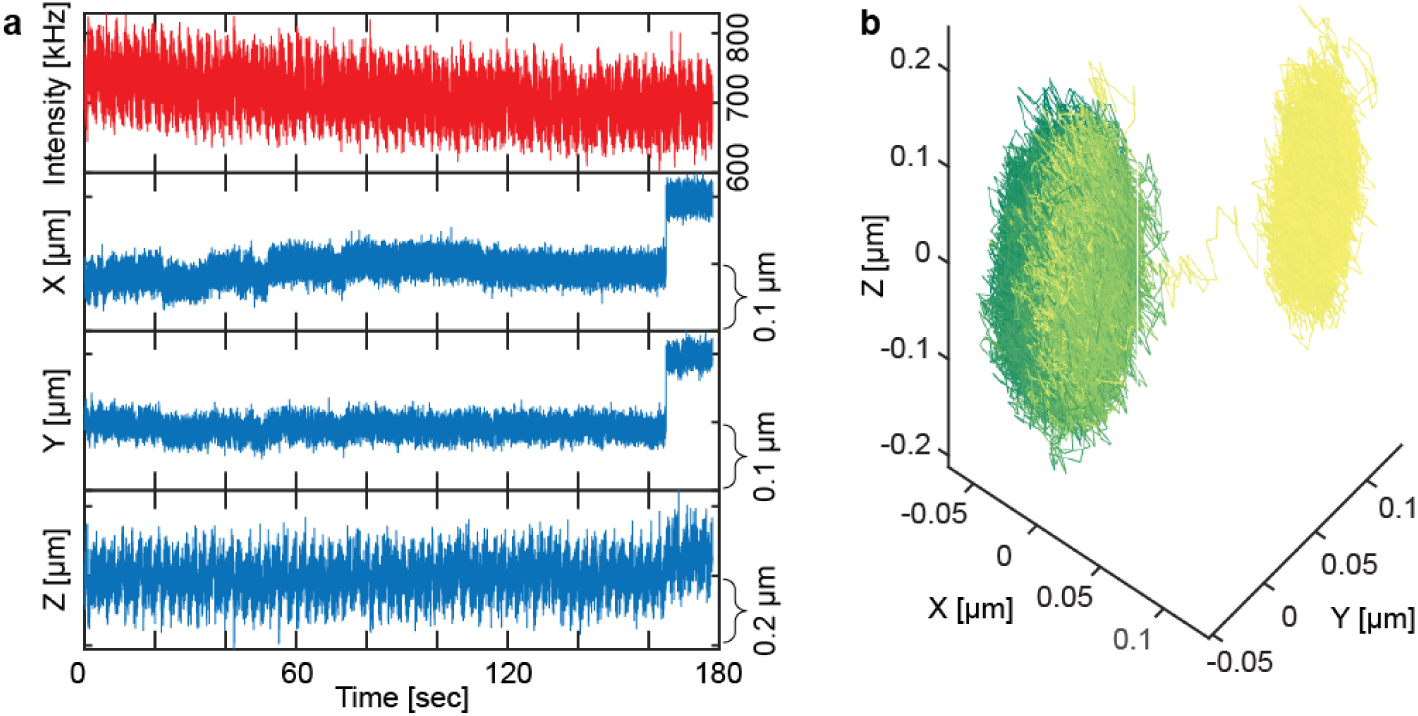
An example of hop events of 500 nm probes in 2 wt% AG. (**a**) Intensity, X, Y, and Z trace. The hop event happened at ∼ 162 sec, and the relocation was less frequent than 200 nm probes in the same concentration of AG. (**b**) An example trajectory of hop events of 500 nm probes in 2 wt% AG.

**Figure S13:**
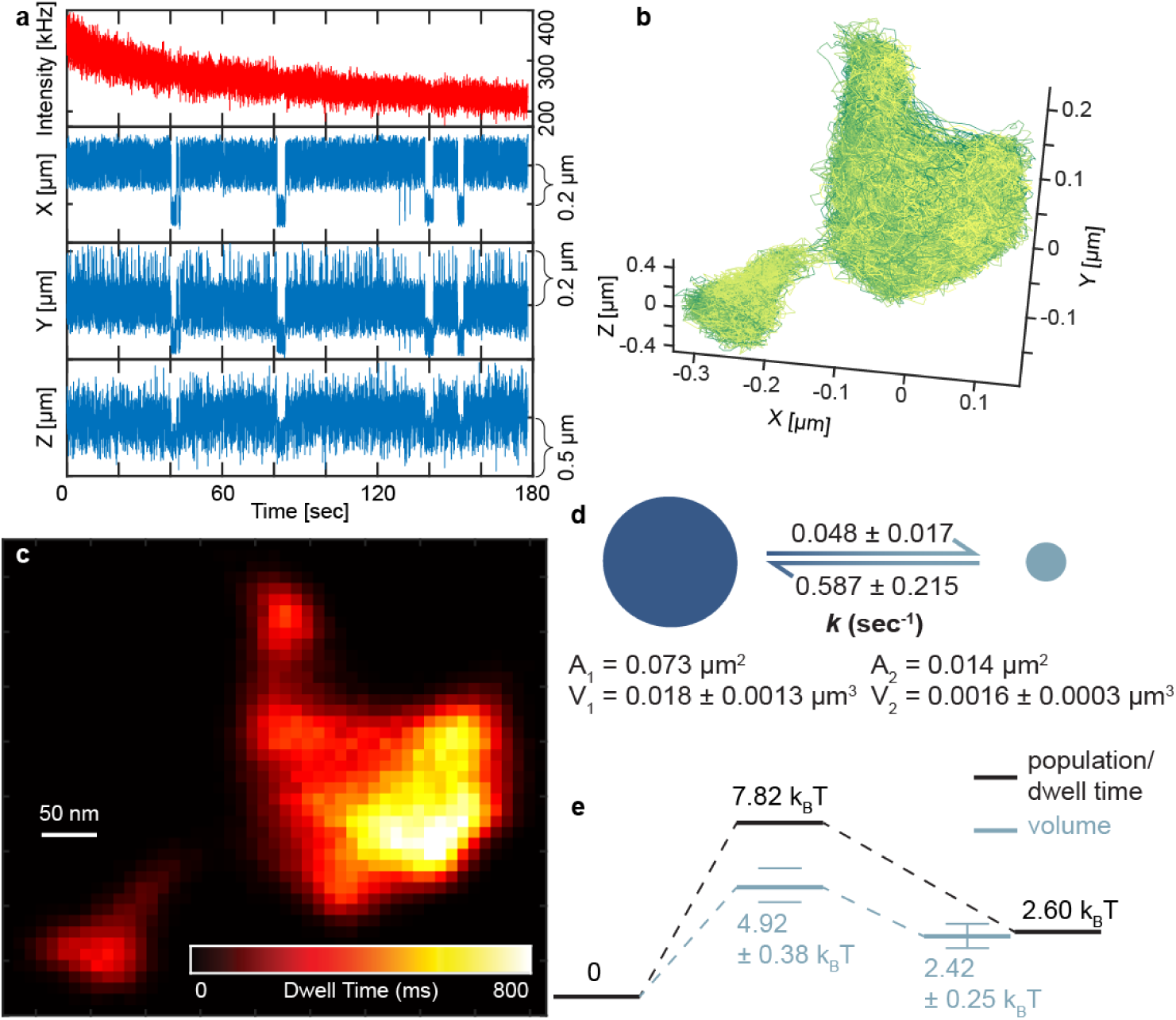
An additional example of hop events with a more static energy barrier. (**a**) Intensity, X, Y, and Z trace. (**b**) An example trajectory of hop events of 200 nm probes in1.5 wt% AG. (**c**) 2D heatmap converted from 3D trajectory based on numbers of localizations (or dwell time) within each pixel (10×10 nm). (**d**) Area, volume, and translocation rate between larger pore 1 and the smaller pore 2. (**e**) Energy barrier between two confinements (calculated based on the population in each pore or based on volume ratio) and activation energy of the hop (calculated based on the population ratio or based on volume ratio). The error bar in free energy barrier (calculated from volume) is determined by expanding or shrinking the channel volume by one localization precision.

**Figure S14:**
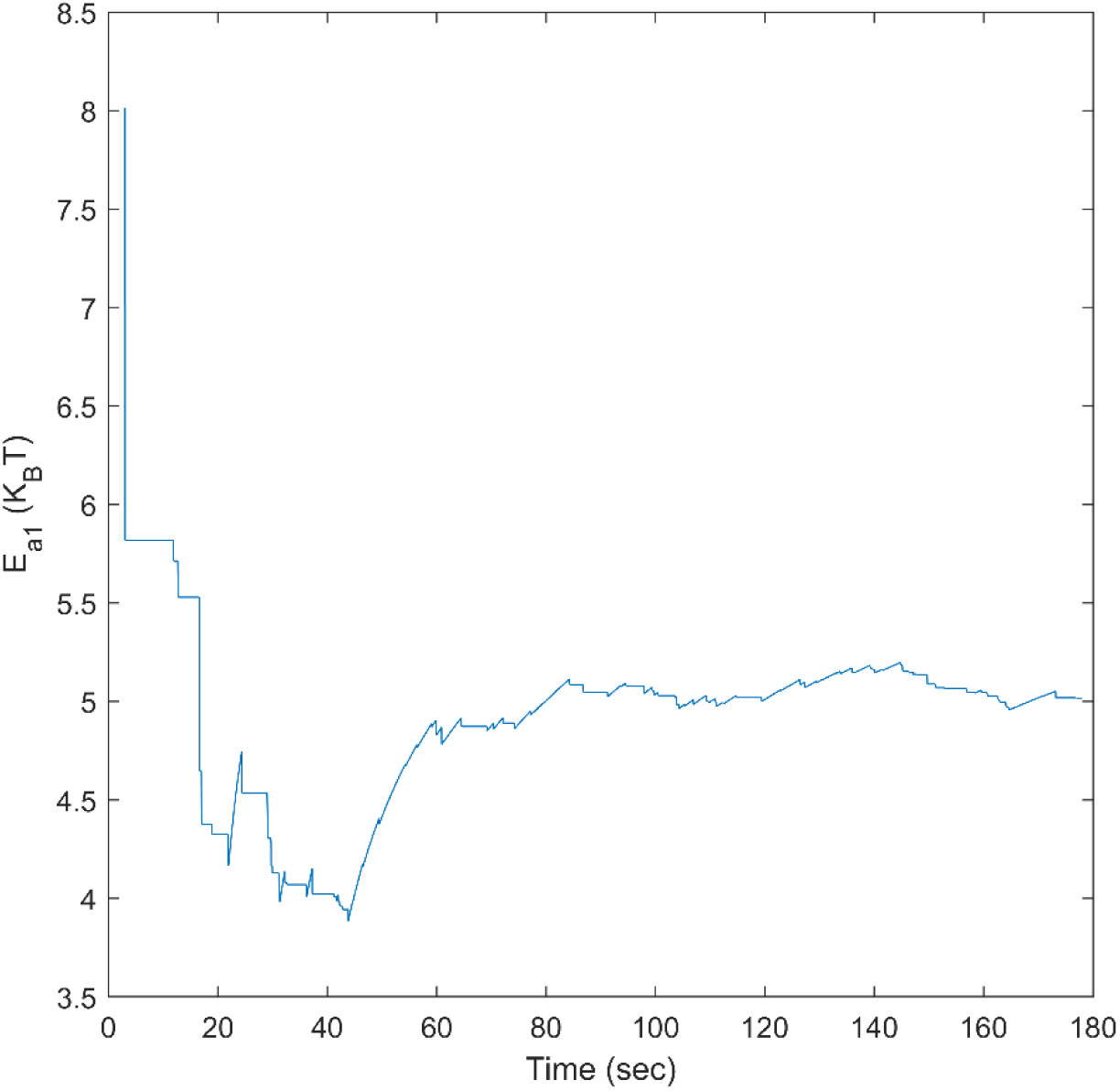
Calculation of energy barrier as a function of time, related to Figure 4. The energy barrier shown in Figure 4 can be calculated as a function of time. It initially fluctuates as a result of kinetic trapping but stabilizes after approximately 1 minute as sampling increases.

**Figure S15:**
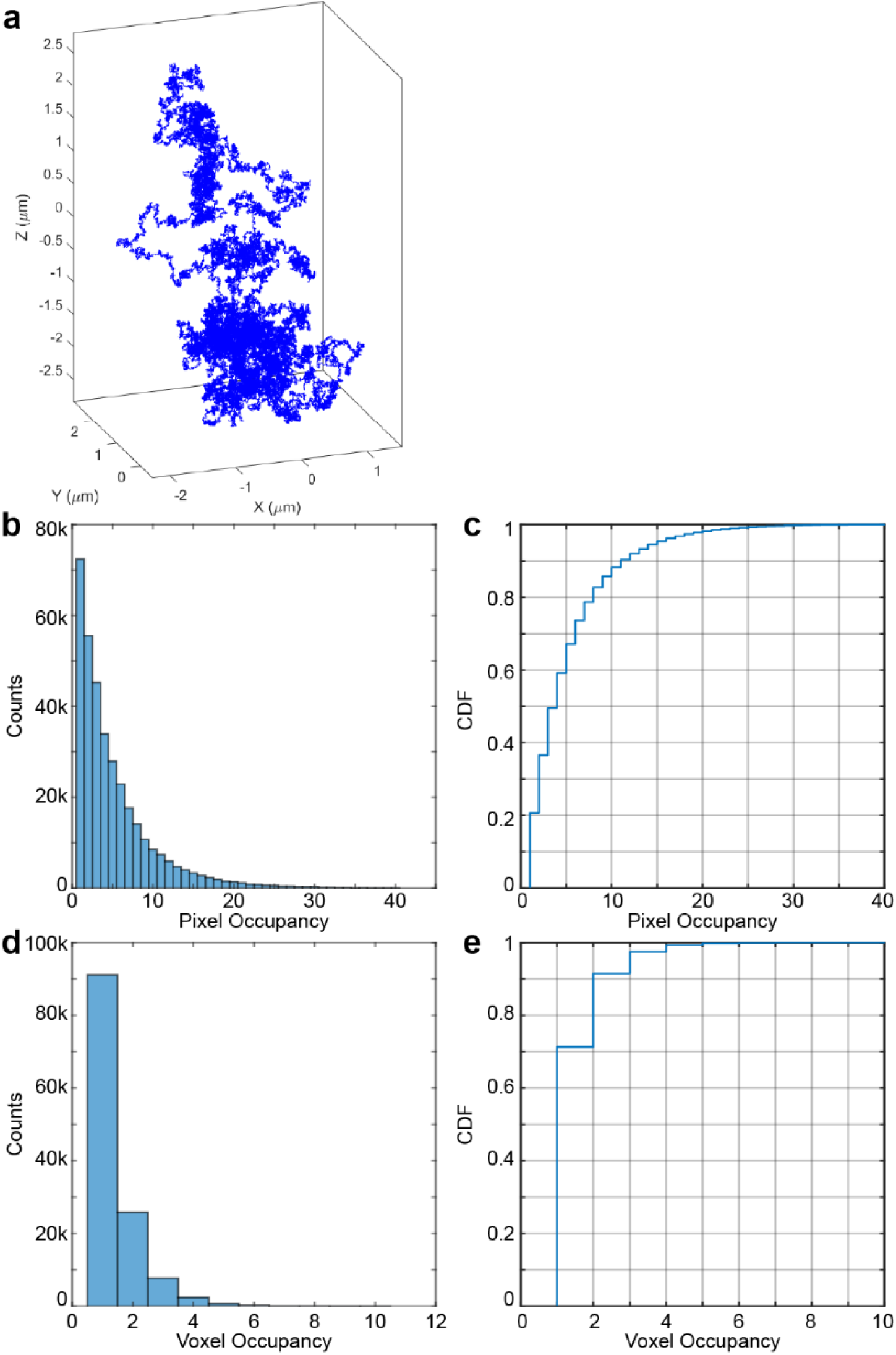
Determining minimum pixel/voxel occupancy through simulated trajectories. (**a**) An example of a 180-sec simulated trajectory with time steps of 1 msec and diffusion coefficient of 0.03 μm2/sec to mimic slow but unconfined motion. (**b**, **c**) Distribution and CDF of pixel occupancy of slow but unconfined diffusion. 15 is chosen as the minimum pixel occupancy for confinement with 95% confidence. (**d**, **e**) Distribution and CDF of voxel occupancy of slow but unconfined diffusion. 2 is chosen as the minimum voxel occupancy for confinement with 92% confidence.

**Movie S1. Hopping diffusion of a 200 nm probe in 2.0 wt% AG, related to Figure 2C-D**.

Left: 3D trajectory (∼ 5 minutes) was color mapped by time with a sampling rate of 1 kHz. Right: X, Y, Z, and intensity traces were synchronized with the 3D trajectory. Playback rate: 10×.

**Movie S2. Hopping diffusion of a 200 nm probe in 2.0 wt% AG, related to Figure 2E**.

3D trajectory (∼ 5 minutes) was color mapped by time with a sampling rate of 1 kHz. Playback rate: 10×.

**Movie S3. Hop events between two adjacent confinements, related to Figure 4**.

(**A**) 3D trajectory (∼ 3 minutes) was color mapped by time with a sampling rate of 1 kHz. (**B**) Photon arrival image time series synchronized with trajectory. 25 grids were color-coded based on the number of photons detected within a bin time, which was 200 ms. Playback rate: 10×. (**C**) 3D point clouds of localizations in two pores (dark blue and light blue) and channel (red).

## Notes

### Competing Interest Statement

The authors have declared no competing interest.

### Summary of Updates

Revisions after peer review, including more information in the introduction, clarification on cavity identification methods, more discussion on the entropic model...

